# X chromosome-dependent disruption of placental regulatory networks in hybrid dwarf hamsters

**DOI:** 10.1101/2020.09.15.298893

**Authors:** Thomas D. Brekke, Emily C. Moore, Shane C. Campbell-Staton, Colin M. Callahan, Zachary A. Cheviron, Jeffrey M. Good

## Abstract

Embryonic development in mammals is highly sensitive to changes in gene expression within the placenta. The placenta is also highly enriched for genes showing parent-of-origin or imprinted expression, which is predicted to evolve rapidly in response to parental conflict. However, little is known about the evolution of placental gene expression, or if divergence of placental gene expression plays an important role in mammalian speciation. We used crosses between two species of dwarf hamsters (*Phodopus sungorus* and *P. campbelli*) to examine the genetic and regulatory underpinnings of severe placental overgrowth in their hybrids. Using quantitative genetic mapping and mitochondrial substitution lines, we show that overgrowth of hybrid placentas was primarily caused by genetic differences on the maternally inherited *P. sungorus* X chromosome. Mitochondrial interactions did not contribute to abnormal hybrid placental development, and there was only weak correspondence between placental disruption and embryonic growth. Genome-wide analyses of placental transcriptomes from the parental species and first and second-generation hybrids revealed a central group of co-expressed X-linked and autosomal genes that were highly enriched for maternally-biased expression. Expression of this gene network was strongly correlated with placental size and showed widespread misexpression dependent on epistatic interactions with X-linked hybrid incompatibilities. Collectively, our results indicate that the X chromosome is likely to play a prominent role in the evolution of placental gene expression and the accumulation of hybrid developmental barriers between mammalian species.

## Introduction

Developing mammalian embryos depend on the extra-embryonic placenta for a broad array of functions including hormone production, immunity, and as a conduit for maternal nutrients and gas exchange (Reik *et al*. 2003; Levy 2007). Normal intrauterine development in humans and mice depends on the tightly controlled placental expression of a diverse set of genes (Constancia *et al*. 2002; Levy 2007; Plasschaert and Bartolomei 2014). Placental gene expression has also likely played an important role in the evolution of mammalian development (Haig 1996; Capellini *et al*. 2011; Kaneko-Ishino and Ishino 2019). Indeed, much of the phenotypic diversity across mammalian species is thought to have evolved by changes in gene expression during critical stages of development (King and Wilson 1975; Carroll 2008; Sears *et al*. 2015). However, relatively little is known about the broader genomic organization, functional integration, and evolution of placental gene expression networks across species (Al Adhami *et al*. 2015).

The placenta is characterized by two unusual regulatory phenomena that likely play critical roles in its evolution. First, the placenta is highly enriched for genes showing monoallelic expression due to epigenetic silencing of one parental allele (i.e., genomic imprinting, Morison *et al*. 2005; Hudson *et al*. 2010; Babak *et al*. 2015). Genomic imprinting is thought to have evolved to help resolve fitness conflicts between maternally and paternally-inherited alleles (i.e., kinship or parental conflict theory, Haig 2000). While perhaps only ∼100-200 autosomal genes show strongly imprinted expression across tissues (Babak *et al*. 2015), disruption of genomic imprinting has emerged as an important cause of congenital disorders in humans (Hirasawa and Feil 2010; Lee and Bartolomei 2013) and as a potential driver of reproductive barriers between species (Crespi and Nosil 2013; Wolf and Brandvain 2014). Second, some mammals also show imprinted paternal X chromosome inactivation in extra-embryonic tissues (i.e., imprinted XCI, Heard and Disteche 2006), representing a striking deviation from random XCI found in most somatic cells of placental mammals (Lyon 1961; Dupont and Gribnau 2013). The X chromosome in general, and imprinted XCI in particular, has been shown to play important roles in placental development (Hemberger 2002; Mcgraw *et al*. 2013). Moreover, the mouse X chromosome appears enriched for genes preferentially expressed in the placenta (Khil *et al*. 2004).

Resolution of genetic conflict may explain the origin of genomic imprinting in mammals (Haig 2000). However, theory also predicts that once imprinting is established adaptation among interacting genes can drive the evolutionary expansion of regulatory networks of similarly imprinted genes (Wolf and Hager 2006; Wolf and Brandvain 2014; Patten *et al*. 2016; O’Brien and Wolf 2017). Given the relative scarcity of autosomal imprinting overall (Babak *et al*. 2015), the X chromosome is expected to harbor the majority of genes showing imprinted (maternal) placental expression in species with imprinted XCI. Thus, co-evolutionary interactions between the maternal X chromosome and maternally expressed autosomal genes should be relatively common within placental regulatory pathways in imprinted XCI species. Despite these predictions, the overall importance of the X chromosome to the evolution of placental gene expression remains unclear. Many molecular genetic studies on the placenta have focused on established disease models or targeted genetic manipulations of imprinted autosomal genes (e.g., gene knockdowns or knockouts, Sanli and Feil 2015), revealing fundamental insights into the mechanisms and functional consequences of genomic imprinting in the placenta and other tissues. In parallel, meta-analyses of expression data have revealed that clusters of imprinted autosomal genes appear to fall within larger networks of co-expressed genes that include both imprinted and bi-allelically expressed loci (Varrault *et al*. 2006; Al Adhami *et al*. 2015). The extent and functional relevance of such regulatory networks remain unclear, but the emerging model of genome-wide networks of imprinted and non-imprinted genes represents a conceptual shift from the view of imprinting controlled primarily through local cis-regulatory effects (Patten *et al*. 2016).

Parent-of-origin effects for abnormal embryonic and placental growth are common in mammalian hybrids (Vrana 2007; Brekke and Good 2014), suggesting that hybrid systems may provide powerful models for understanding how the evolution of gene expression impacts placental development. Here, we focus on crosses between two closely related species of dwarf hamsters (*Phodopus* sungorus and *P. campbelli*) that yield strong parent-of-origin growth effects in reciprocal F_1_ crosses (Brekke and Good 2014). Extensive placental overgrowth and disrupted embryonic development occur in hybrid crosses when the mother is *P. sungorus* (female *P. sungorus* x male *P. campbelli*; hereafter S×C with uppercase used to denote placental overgrowth), often resulting in hybrid and maternal death during birth. The reciprocal cross results in normal embryonic development (female *P. campbelli* x male *P. sungorus*; hereafter c×s with lowercase used to denote normal placental growth), although adult hybrid males are smaller (Brekke and Good 2014) and completely sterile (Safronova *et al*. 1999; Bikchurina *et al*. 2018). Intrinsic postzygotic reproductive isolation (i.e., hybrid inviability or sterility) generally tends to be asymmetric in reciprocal crosses between closely related species due to incompatible interactions at sex-linked or imprinted loci (Turelli and Moyle 2007). Although the genetic architecture of hybrid placental growth has not been determined in dwarf hamsters, massively overgrown F_1_ hybrid S×C placenta do show extensive disruption of gene expression pathways that are highly enriched for embryonic growth and genomic imprinting (Brekke *et al*. 2016). Building on these previous studies, we combine quantitative genetic and transcriptomic analyses to test the hypothesis that the X chromosome plays a central role in the evolution of placental gene expression, embryonic development, and reproductive barriers between species.

## Materials and Methods

### Animals and experimental crosses

Experimental animals were drawn from established colonies of wild-derived *P. campbelli* and *P. sungorus* at the University of Montana (Brekke and Good 2014), which were originally established by Kathy Wynne-Edwards (Scribner and Wynne-Edwards 1994). Colonies were maintained as outbred, though overall inbreeding levels are high (Brekke *et al*. 2018). All breeding experiments were done in compliance with the University of Montana Institutional Animal Care and Use Committee regulations (animal use protocols 039-13JGDBS-090413 & 050-16JGDBS-082316).

We previously reported results from experimental crosses within and between *P. campbelli* and *P. sungorus* used to examine late-term placental and embryonic phenotypes (Brekke and Good 2014) and placental transcriptomes (n=40 placental transcriptomes, 5 males and 5 females for each cross type; Brekke *et al*. 2016). Here we combined these results with new data from two additional genetic crossing experiments used to evaluate the role of the mitochondrial and nuclear genomes in the genetic basis of asymmetric hybrid placental and embryonic overgrowth (Figure 1). First, we generated mitochondrial substitution lines wherein c×s hybrid females were crossed to *P. sungorus* males for ten additional backcross generations. This crossing scheme is predicted to recover hamsters that are ∼99.9% *P. sungorus* across the nuclear genome but retain the mitochondria of *P. campbelli* (*P. sungorus*^mtC^). Tenth-generation *P. sungorus*^mtC^ females were crossed to *P campbelli* males to test for F_1_ overgrowth (S^mtC^ ×C), thereby mimicking the overgrown S×C hybrid across the nuclear genome while substituting the *P. sungorus* mitochondria for *P. campbelli* mitochondria. Second, we crossed F_1_c×s hybrid females to *P. campbelli* males to generate a backcross (BC) panel of late-term embryos and placentas. In the context of genetic elements with sex-limited inheritance or expression, these backcross hybrids have the same paternal contribution found in overgrown S×C F_1_ hybrids, while varying the species origin of maternally inherited alleles.

**Figure 1.**
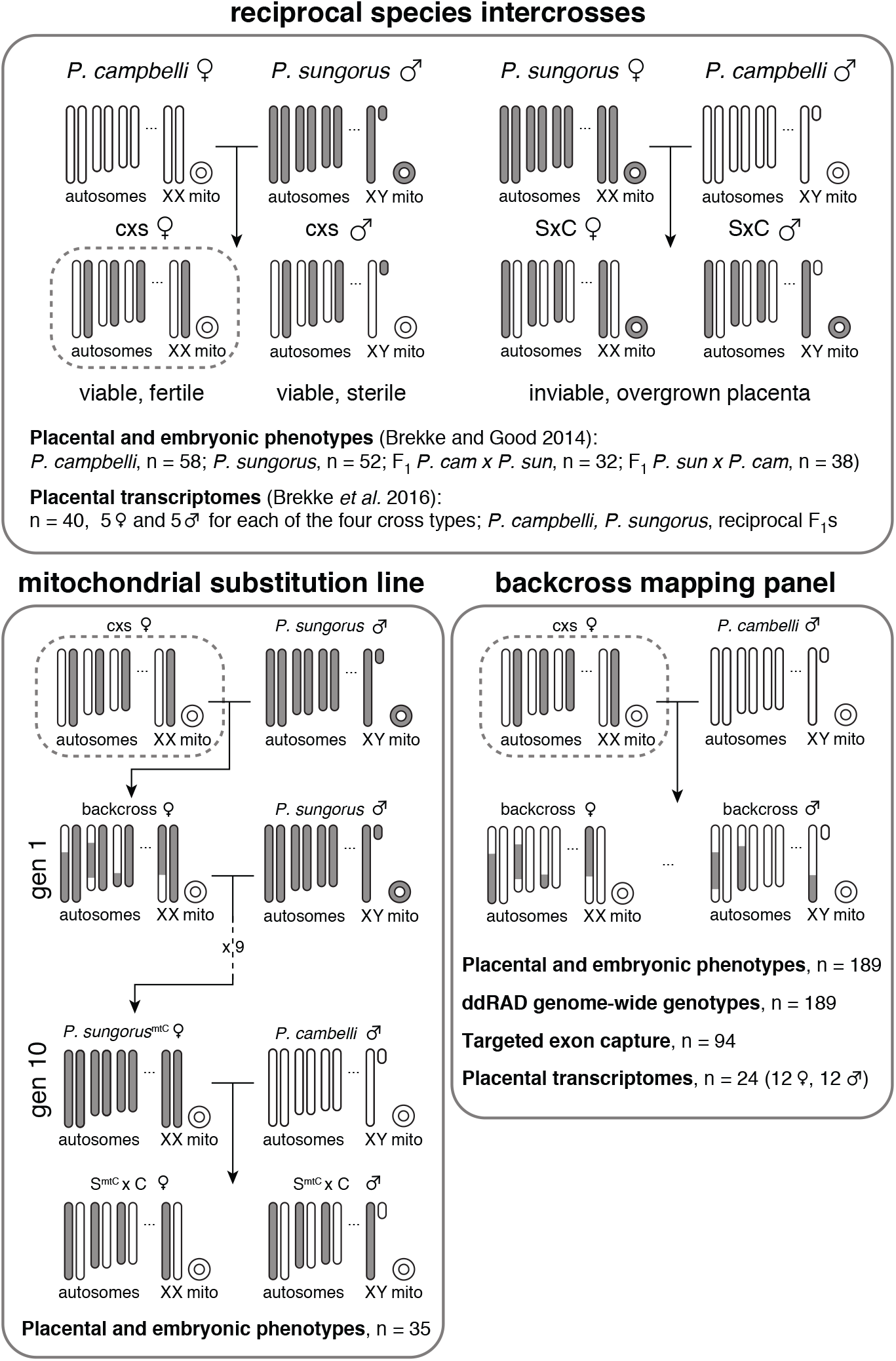
Summary of genetic crosses and experiments. Reciprocal species intercrosses between *P. sungorus* (gray chromosomes) and *P. campbelli* (white chromosomes) result in asymmetric placental overgrowth and inviability. Only females with a *P. campbelli* mother and *P. sungorus* father are both viable and fertile (c×s, indicated with a dashed box). The first generation cross with normal sized placentas is indicated with lowercase letters (cxs), and the reciprocal cross with overgrown placentas is indicated with uppercase letters (SxC). A mitochondrial substitution line was created through ten generations of backcrossing hybrid females to a *P. sungorus* male, resulting in females with *P. sungorus* nuclear genomes and *P. campbelli* mitochondria (S^mtC^) that were then crossed to *P. campbelli* males. The backcross mapping panel was created by a single generation of backcrossing a fertile hybrid female to a *P. campbelli* male, resulting in offspring showing a range of placenta phenotypes. Data types collected for each experiment are reported in the corresponding panels.

All advanced crosses were conducted through c×s hybrid females as S×C F_1_ hybrids generally do not survive birth (Brekke and Good 2014) and c×s males are completely sterile (Safronova *et al*. 1999; Ishishita *et al*. 2015; Bikchurina *et al*. 2018). For both crossing experiments, females were sacrificed at late gestation and offspring placentas and embryos were collected, weighed, and snap-frozen on dry ice. Embryos were developmentally scored following (Brekke and Good 2014) to ensure that all offspring were in the final four days of gestation corresponding to Theiler’s Stages 24-27 (Theiler 1972). Developmental abnormalities such as embryonic fluid accumulation (edema) and embryo reabsorption were noted, and embryo and placenta weights were assessed with stepwise model selection and adjusted for variation in edema, litter size and Theiler stage using simple linear models as implemented in JMP (v12).

### Genotyping

We genotyped 189 backcross individuals (91 females and 98 males) and our original colony founders (14 *P. campbelli* and 11 *P. sungorus*) using double digest restriction-site associated DNA sequencing (ddRAD-seq; Peterson *et al*. 2012) with minor modifications. Genomic DNA was extracted from frozen embryos with a Machery-Nagel Nucleospin Tissue DNA extraction kit (catalog number 740952) following the manufacturer’s protocol, except that 5µl RNase-A was added to the column and incubated for 15 minutes at room temperature. We then digested 1µg of genomic DNA using the high-fidelity restriction enzyme SbfI-HF (New England BioLabs, catalog number R3642L), followed by MspI (New England BioLabs, catalog number R0106L) both with the CutSmart buffer (New England BioLabs). Libraries were prepared with a dual barcoding scheme incorporating both Illumina indexes and in-line barcodes to uniquely identify each sample (Peterson *et al*. 2012). Size selection of adapter-ligated fragments (200-500bp) was done with Agencourt AMPure XP beads. Both the size selection and PCR amplification were done prior to sample pooling to assure more even representation across samples. Combined sample pools were sequenced on 50% of an Illumina HiSeq 2500 lane in rapid-run mode and then on 50% of a lane of Illumina Hiseq 2500 in normal mode at the University of Oregon Genomics and Cell Characterization Core Facility. All samples were sequenced in both lanes and combined for subsequent analyses. We also independently determined the sex of embryos using a PCR assay of the Y-linked Sry gene as described previously (Brekke *et al*. 2016).

RAD libraries were cleaned and demultiplexed with Stacks (v1.20; process_radtags parameters -e sbfI --renz_2 mspI –r -c -q; Catchen *et al*. 2013). An initial list of unique RADtags from both read pairs was generated using ustacks (−H -r-d) using data from one female founder from each species. RADtag reference libraries were then generated using cstacks (-n 4). Reads from all the colony founders were aligned to the RADtag reference library with bwa-mem (v0.7.9a; Li and Durbin 2009) and single-nucleotide variants (SNVs) were called with the GATK HaplotypeCaller (v3.1-1, -stand_call_conf 30; Van Der Auwera *et al*. 2013). All SNVs that were polymorphic within a species in our colony founders were filtered out using GATK selectVariants (v3.1-1). Backcross individuals were genotyped at the ascertained SNVs sites using GATK UnifiedGenotyper (v3.1-1; -stand_ call_conf 30; Van Der Auwera *et al*. 2013).

We also used an exon capture experiment to anchor placental-expressed genes onto the *Phodopus* genetic map (described below). We designed custom hybridization probes to target 9,756 fixed SNVs between *P. campbelli* and *P. sungorus* ascertained from species-specific transcriptomes (Genbank BioProject PRJNA306772 and DDBJ/EMBL/GenBank Accessions GEVA00000000 and GEVB00000000; Brekke *et al*. 2016). Exon boundaries were annotated for each transcript with a local BLAT search (Kent 2002) against the golden hamster (*Mesocricetus auratus*) draft reference genome (The Broad Institute Genome Assembly & Analysis Group, MesAur1.0). For each gene, we selected 1-2 SNVs located furthest from inferred exon boundaries and included probes matching both alternative bases to avoid species bias. Capture baits were manufactured by MYcroarray.

We selected 94 individuals (44 males and 50 females) from the backcross panel and prepared Illumina sequencing libraries following the Meyer-Kircher protocol (Meyer and Kircher 2010). Ten cycles were used during the indexing PCR step. The indexed libraries were then combined into four pools and target enriched following the MyBaits-1 Custom Target Capture protocol and using mouse CoT-1 DNA supplied by the manufacturer as a blocking agent. The four captured pools were then reamplified with 20 PCR cycles, quantified with a Kappa qPCR quantification kit (catalog number KK4824), and pooled for Illumina sequencing. Enriched libraries were initially sequenced at the University of Montana Genomics Core on an Illumina MiSeq 75bp paired-end sequencing, and followed by one lane of Illumina HiSeq 2500 100bp single-end sequencing at the University of Oregon Genomics and Cell Characterization Core Facility. Raw sequence reads were adapter trimmed with Cutadapt (v1.6; -O 5 and -e 0.1; Martin 2011) and quality filtered with Trimmomatic (v0.3.2; LEADING:5, SLIDINGWINDOW:4:15, MINLEN:36, and HEADCROP:13; Bolger *et al*. 2014). Filtered reads were then aligned to published transcriptome assemblies (Brekke *et al*. 2016) and the targeted SNVs were genotyped with GATK HaplotypeCaller (v3.1-1; -stand_call_conf 30.0 -stand_emit_conf 30) and filtered (selectvariants--restrictAllelesTo BIALLELIC -select “QD > 10.0”) so that only high-quality genotypes were used for estimating the location of each gene.

### Quantitative genetic analysis

We constructed a genetic map using RADtag SNVs identified between the *P. campbelli* and *P. sungorus* colony founders and the program R/qtl (v1.45; Broman and Sen 2009). Putative X-linked RADtags were identified as markers that were heterozygous or homozygous for *P. campbelli* genotypes in females and always homozygous *P. campbelli* or *P. sungorus* in males. To estimate the map, we removed two backcross individuals with low sequencing coverage, identified putative X-linked markers based on Hardy-Weinberg expectations, and dropped all autosomal markers that were genotyped in fewer than 177 individuals (95%). We formed linkage groups and ordered the markers on each linkage group with the ripple(), compareorder(), and switch.order() functions until each linkage group was a short as possible. Then we sequentially dropped each marker to see if the likelihood of the map improved. Once all poor quality markers were removed, we repeated the ripple(), compareorder(), and switch.order() functions until the likelihood was maximized. The linkage groups in the final map were ordered by descending length in centiMorgans (cM).

We then used R/qtl to test for single quantitative trait loci (QTL) associated with the variation in backcross embryo and placental weight using the extended Haley-Knott method with imputation (Haley and Knott 1992; Feenstra *et al*. 2006). Next, we incorporated sex as a covariate and re-estimated QTL for both phenotypes. For all of single-QTL scans, we used 10,000 permutations to estimate genome-wide significance thresholds for autosomes and 337,364 permutations for the X chromosome. Finally, we used the QTL identified in the first two analyses as additive cofactors and re-scanned for additional QTL that were contingent on the presence of the earlier identified QTL (Broman and Sen 2009). We used 1,000 permutations for autosome-autosome interactions, 1,687 permutations for autosome-X interactions, and 113,815 permutations for X-X interactions to establish significance thresholds. QTL intervals were established based on 95% Bayesian confidence intervals, and the proportion of phenotypic variance explained by QTL was estimated as 1-10^(-2/n^ ^LOD)^ (Broman and Sen 2009). Expressed genes were then integrated onto the genetic map by comparing captured SNV genotypes to RADtag genotypes. Following Brekke *et al*. (2019), we counted shared genotypes between each RADtag and each gene across all individuals, placing genes at the location of the RADtag with which they shared the most genotypes. In the event of a tie between multiple RADtags, the gene was placed at the proximal map location and only genes sharing at least 90% of genotypes with at least one RADtag were placed on the map. Given the low number of recombination events and the high number of genes, these RADtag-anchored positions represent coarse genetic locations for each gene. Instances where multiple genes were associated with a single RADtag were treated as a single unordered linkage block. Once integrated, likely genotyping errors in the capture data were identified using calc.errorlod() and the highest errors were extracted with top.errorlod() and a very strict cutoff of 1 such that even moderately questionable genotypes were identified. The genotypes of these sites were removed and then filled in with the fill.geno() function using the imputation method, which imputes the missing genotypes from the surrounding sites for each individual. These corrected genotypes were used for evaluating imprinting status of select candidate genes (see below).

### Gene expression analyses

We chose 24 placentas from our backcross mapping panel for genome-wide expression analysis using RNA-seq (Wang *et al*. 2009), including six males and six females with large placentas (mean = 0.232±0.010g) and six males and six females with normal sized placentas (mean = 0.140±0.008g). RNA was extracted from whole frozen placenta with an Omega Bio-tek E.Z.N.A. Total RNA Kit I (catalog number R6834) including a DNase digestion following the manufacturer’s protocol. All RNA samples were checked for quality and concentration on the bioanalyzer and all samples used had RNA integrity numbers (RIN) greater than 8.0. RNA-seq libraries were constructed from 2 µg of input RNA with the Agilent Sure-Select Strand-Specific RNA-seq Kit (catalog number G9691B) following the manufacturer’s recommendations. Libraries were amplified with 14 cycles of PCR, and pooled based on a Kappa qPCR Quantification Kit (catalog number KK4824). The pooled libraries were sequenced with two lanes of Illumina HiSeq2500 100bp single-end sequencing.

RNA-seq data were processed as described previously (Brekke *et al*. 2016). Briefly, Illumina adapters were trimmed off reads with Cutadapt (v1.6; -O 5 -e 0.1; Martin 2011) and quality trimmed with Trimmomatic (v0.3.2; SE -phred 33 LEADING:5 SLIDINGWINDOW:4:15 HEADCROP:13; Bolger *et al*. 2014). While an initial draft of the *P. sungorus* genome has been generated using Illumina shotgun sequencing, current annotation and assembly quality remains insufficient for reference-guided transcriptome analyses (Bao *et al*. 2019). Therefore, reads were aligned with bowtie2 (v2.2.3; Langmead and Salzberg 2012) to a published de novo placental transcriptome assembly (Genbank BioProject PRJNA306772 and DDBJ/EMBL/ GenBank Accessions GEVA00000000 and GEVB00000000; Brekke *et al*. 2016), and filtered for potentially chimeric transcripts using draft genomic resources by excluding 1,422 ‘genes’ with exons that multiply mapped to different contigs. To evaluate expression level, we created a table of counts at the gene level using featureCounts (v1.4.2; Liao *et al*. 2014), which counted fragments (-p) and discarded reads with too long an insert (-P) or are chimeric (-C) or have a mapping quality (-Q) below 20. This table of counts was normalized with the TMM method (Robinson and Oshlack 2010).

We used the WGCNA package (version 1.68; Langfelder and Horvath 2008) to infer weighted gene co-expression networks using expression data from previously published parental and F_1_ genotypes (Brekke *et al*. 2016) and the newly generated backcross. This network approach uses adjacency correlation matrices to identify hierarchical clusters of co-expressed genes (Zhang and Horvath 2005), enabling the reduction of complex clusters into representative expression profiles (i.e., module eigengenes) defined as the first component of a principle component analysis. A scale-free topology index was determined by soft thresholding (Figure S1), which was then used to automatically detect signed, Pearson correlated modules via dynamic cutting. The signed module method splits otherwise correlated genes with increases in expression into separate modules from those with decreases in expression, which allowed us to evaluate upregulated gene sets independently from downregulated gene sets. We merged similar modules using a threshold of 0.25, which combines modules with a correlation of 0.75 or greater into a single module. For each module, we tested for correlations between the module eigengene and placental and embryonic weights. For modules showing significant correlations after correction for multiple testing, we then retested associations using a more stringent ANOVA model that controlled for developmental stage and sex.

Each module was assessed with a binomial exact test (R/ stats package 3.6.1) for enrichment of candidate imprinted genes previously identified based on patterns of allele-specific expression in the placenta (Brekke *et al*. 2016) and for X-linked genes. Network connectivity was determined through pairwise correlation between genes, with p-values corrected with the qvalue package (version 2.18.; Storey *et al*. 2019), using a false discovery rate threshold of 0.05. We then counted the number of additional genes significantly correlated with each gene within the module. Hub genes were defined as the top 5% most connected genes in each module. Overlap between F_1_ and backcross modules was determined by comparing gene lists to get counts of shared genes. We also evaluated module conservation by comparing how strongly each gene was correlated to each module across data sets. To connect module conservation to phenotypes, we compared the concordance between each gene and placenta weight across F_1_ and backcross datasets using a bivariate correlation as implemented in JMP (v12).

Genotypes at each gene placed on the genetic map were inferred by evaluating genotypes at flanking RAD markers. If a gene was placed in the same linkage block (cM location) as a RADtag, the marker genotype was used for the gene. Likewise, if the gene was placed between two markers with the same genotype, the concordant marker genotypes were used for the gene. If flanking RADtag genotypes were discordant, the genotype at the gene was treated as unknown. We then tested for expression interactions between the X chromosome and diploid autosomal genotypes within the backcross as follows. Autosomal genotypes were imputed for the 23 BC individuals with transcriptome data. Expression interaction (EI) scores were then calculated for all mapped autosomal genes by comparing the difference in fold change in expression of that gene for the two possible autosomal genotypes (i.e., homozygous for the *P. campbelli* allele or heterozygous) dependent on the genotype of the maternal X chromosome (*P. campbelli* or *P. sungorus*). To generate EI scores for each gene, normalized gene expression count tables generated during WGCNA were used to calculate log2-fold changes between alternative maternal X chromosome genotypes and the two genotypes at each autosomal locus. We excluded unmapped autosomal genes and genes with imputed genotypes for fewer than three individuals for each of the four X-by-autosome genotypic classes. We calculated the absolute value of EI scores, where a value of zero indicates no difference between the two autosomal genotypes when inheriting different maternal X chromosomes and a value of one indicates a one-fold difference in expression between the two autosomal genotypes (i.e., a candidate X-autosome expression interaction). We also considered a polarized (i.e., non-absolute value) version of the statistic where positive values reflected greater change when maternal X and autosomal alleles genotypes were discordant.

EI was calculated as follows:

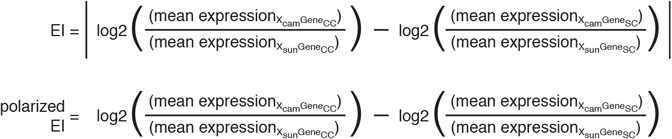

where:

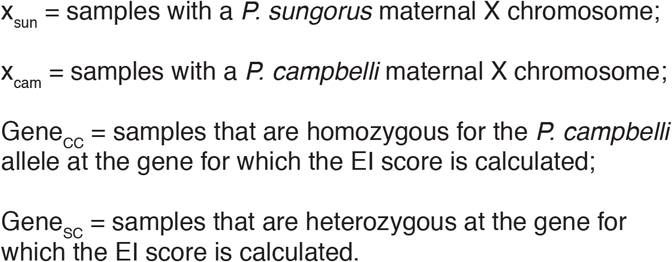

### Data Availability

All raw sequencing reads are archived at NCBI under BioProject PRJNA306772. Accession numbers for individual libraries are provided in Table S5. All other data are provided in Data S1 as supplementary material available at Figshare: https://doi.org/10.6084/m9.figshare.13664603.v3

## Results

### Mitochondrial interactions were not a major factor in extreme parent-of-origin hybrid overgrowth

If hybrid placental overgrowth is caused by deleterious interactions between the maternal *P. sungorus* mitochondrial genome and the *P. campbelli* nuclear genome, then crossing females from the *P. sungorus*^mtC^ mitochondrial substitution lines to *P. campbelli* males should recover normal development. Alternatively, if heterospecific mitochondria have no effect on hybrid growth, then S^mtC^×C hybrids should be similar in size to overgrown S×C hybrids. In support of the second hypothesis, S^mtC^×C placentas were found to be extremely large (F_4,213_ = 106, P<0.001, ANOVA), but not statistically different from S×C hybrids based on a Tukey’s HSD test (Figure S2). We note that low levels of residual autosomal heterozygosity could theoretically mask mitochondrial incompatibilities in the tenth generation mitochondrial substitution *P. sungorus*^mtC^ lines. However, such residual variation cannot also explain why the substitution lines did not recover normal placental development in the F_1_ hybrid crosses. On balance, these results strongly indicate that heterospecific mitochondrial interactions are not a major factor in extreme parent-of-origin hybrid overgrowth in dwarf hamsters.

### The quantitative genetic basis of extreme parent-of-origin hybrid overgrowth

We generated a backcross panel of 189 late-term embryos and placentas sampled from 38 litters. Using 1,215 ddRAD SNVs, we constructed a 1,231.7 cM genetic map comprised of 14 major linkage groups (Figure S3, Table S1). The karyotype of *P. sungorus* is comprised of 14 chromosomes (2N=28; Romanenko *et al*. 2007), suggesting that each of our major linkage groups correspond to an individual chromosome. The X chromosome was inferred to have the shortest overall genetic distance (35.5 cM or 2.9% of the genetic map), while it appears medium-sized in ranked karyotype analyses (i.e., middle 30%; Gamperl *et al*. 1977; Romanenko *et al*. 2007) and comprises ∼10% of the haploid female karyotype (Haaf *et al*. 1987). The *Phodopus* X chromosome is metacentric with an Xp arm that has been described as heterochromatic (Gamperl *et al*. 1977) and non-recombinant in females of both species and in c×s F1 hybrids (Bikchurina *et al*. 2018). Our inferred genetic map was consistent with strong repression of recombination on one end of the X chromosome in c×s females, albeit not complete repression as suggested in recent study that quantified signals of mismatch repair through immunolocalization of *MLH1* (Bikchurina *et al*. 2018). Using this genetic map, we then tested for QTL associated with variation in backcross embryo and placenta weights (Figure 2). We observed a single QTL of large effect for placental weight on the X chromosome, with 52.3% of the observed variation in backcross placental weight determined by the genotype of the maternally inherited X chromosome (Figure 2C). We estimated a QTL peak at 31.1cM with a 95% Bayesian confidence interval between 29.6cM and 32.6cM. This X-linked QTL localized near the proximal boundary of where we also observed repressed recombination (Figure S4), although the entire X chromosome exceeded a permutation-based significance threshold (P = 0.01). Male and female embryos inheriting a maternal *P. sungorus* X chromosome genotype at this QTL showed an ∼60% increase in average placenta weight (F_1,179_ = 178.4, P < 0.0001, ANOVA; Figure 2C inset). No additional placental QTL were uncovered when using sex, developmental stage, and litter size as covariates, nor when using the X-linked QTL as a cofactor.

**Figure 2.**
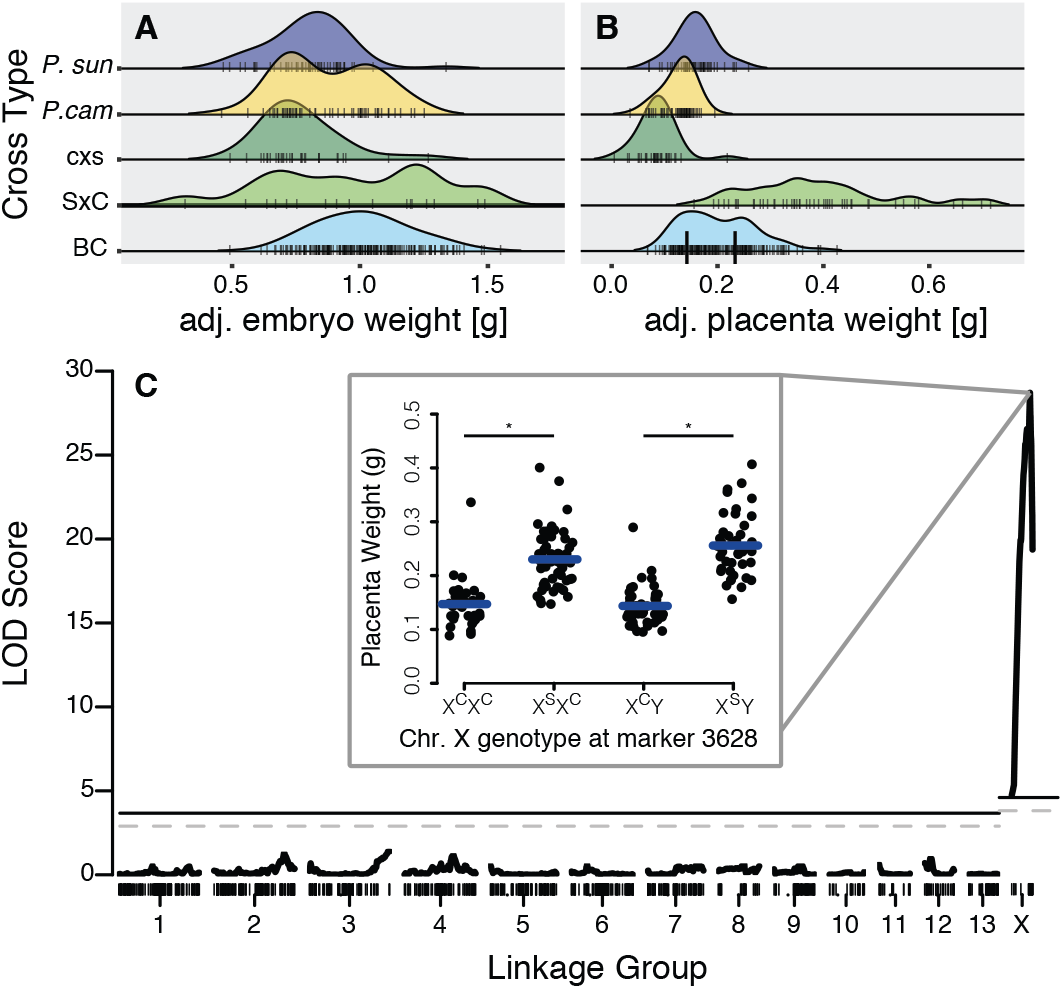
Phenotype Distributions and QTL for placental size. Late term embryo size adjusted for Theiler stage and edema (A) and placenta size adjusted for Theiler stage (B) for *P. sungorus, P. campbelli*, reciprocal F_1_s and 189 BC conceptuses. (C) Genetic mapping revealed a single QTL for placenta weight on the X chromosome. The *P. sungorus* X chromosome increases placenta weight by ∼60% (inset, F_1,179_ = 178.4, P << 0.0001, stars indicate significant differences assigned by a Tukey’s HSD test). Placenta weights were grouped by the genotype at marker 3628. Genotypes are denoted with the maternal allele first followed by the paternal allele. Significance thresholds are denoted by solid (P = 0.01) and dashed (P = 0.05) horizontal lines.

No QTL for embryo weight were recovered in our experiment (P = 0.05 permutation-based significance threshold; Figure S5), despite considerable variation in backcross embryo weights (Figure 2A) and significant overgrowth of S×C embryos when compared to normally developing cross-types (Brekke and Good 2014). Severe embryonic swelling or edema is common in SxC hybrids and appears to drive overall differences in embryo weights between overgrown SxC hybrids and either species or cxs hybrids (stepwise best model: embryo weight ∼ developmental stage + edema; adjusted r2 = 0.40, F_2,182_ = 61.6, P << 0.0001; adjusted embryo weight ∼ cross type F_3,182_ = 0.74, P = 0.5312, ANOVA; Figure S6). However, backcross placenta and embryo weights remained moderately correlated after adjusting for developmental stage and edema (adjusted r^2^ = 0.159, F_1,184_ = 36.0, P << 0.0001, ANOVA; Figure S7A), with males showing a stronger correlation than females (males: adjusted r^2^ = 0.257, F_1,95_ = 33.8, P << 0.0001; females: adjusted r2 = 0.065, F_1,88_ = 7.15, P = 0.0090, ANOVA). When we expanded our analysis of embryonic weights among genotypes to include the backcross, we also detected a small but significant increase in embryo weight in the overgrown crosses after controlling for age and edema (adjusted r^2^ = 0.176, F _4,376_ = 21.1, P << 0.0001, ANOVA; Figure S7B). These apparent differences in embryonic growth were likely too subtle to detect in our QTL mapping experiment.

### Networks of placental gene expression in first generation reciprocal hybrids

We previously demonstrated extensive parent-of-origin dependent disruption of hybrid gene expression, with hundreds of genes significantly up- or down-regulated in overgrown S×C hybrid placenta relative to both species (i.e., transgressive expression, Brekke *et al*. 2016). In this previous study, examination of allele-specific expression revealed 88 candidate imprinted genes in the placenta overall, with 79 genes showing strong bias towards maternal expression (i.e., paternally imprinted). Notably, 68% of candidate imprinted genes (60 genes) showed transgressive expression in overgrown hybrids, suggesting a link between misexpression of autosomal genes with biased parent-of-origin expression and placental overgrowth. Imprinted XCI was not disrupted in F_1_ hybrids. In contrast to a predominantly autosomal regulatory phenotype in F_1_ hybrid placenta, our backcross experiments indicated that S×C F_1_ hybrid placental overgrowth was likely caused by genetic incompatibilities exposed on the maternally inherited *P. sungorus* X chromosome (Figure 2C).

Motivated by these parallel observations, we sought to identify groups of co-expressed genes associated with overgrowth phenotypes exposed in both of our first and second generation hybrid models of placental overgrowth. We first used our published late-term placental expression data from the parental species and reciprocal F_1_ hybrids to construct weighted gene co-expression networks, removing one female SxC sample during filtering (n=39 placental transcriptomes, 5 males and 5 females per cross-type; Figure S8). We placed 11,392 genes (including 70 candidate imprinted genes) into 29 signed clusters, or ‘modules,’ of non-overlapping gene sets. For each module, expression values were summarized with an ‘eigengene’, or the principal component capturing the largest proportion of the variance in gene expression. We then assessed each module for mode of eigengene inheritance and association with placental phenotypes (Table S2 and Figure S9). Two key gene networks emerged from this analysis. One module was comprised of 565 genes that tended to be highly expressed in S×C hybrid placenta relative to all other genotypes. The eigengene value for this module was positively correlated with placental weights (Figure 3A), and included only one candidate imprinted gene (Figure 3B). The other module was comprised of 1160 genes that tended to show a lower eigengene value in S×C hybrid placenta (Figure 3C). Expression of this downregulated set was negatively correlated with placental weights (Figure 3C), and included nearly half (44%) of the downregulated transgressive genes identified by Brekke *et al*. (2016). Eigengene values for this module exhibited a stronger parent-of-origin mode of inheritance than the upregulated set (Figure 3C), and was positively correlated with candidate imprinted gene expression (Figure 3D). The downregulated module included over 50% of the candidate placental imprinted genes overall (36 of 70 genes, P << 0.0001; Table S2,). These findings mirror results from pairwise contrasts (Brekke *et al*. 2016) where overgrown S×C placentas showed an overall reduction in the expression levels of several putatively imprinted genes (Figure 4).

**Figure 3.**
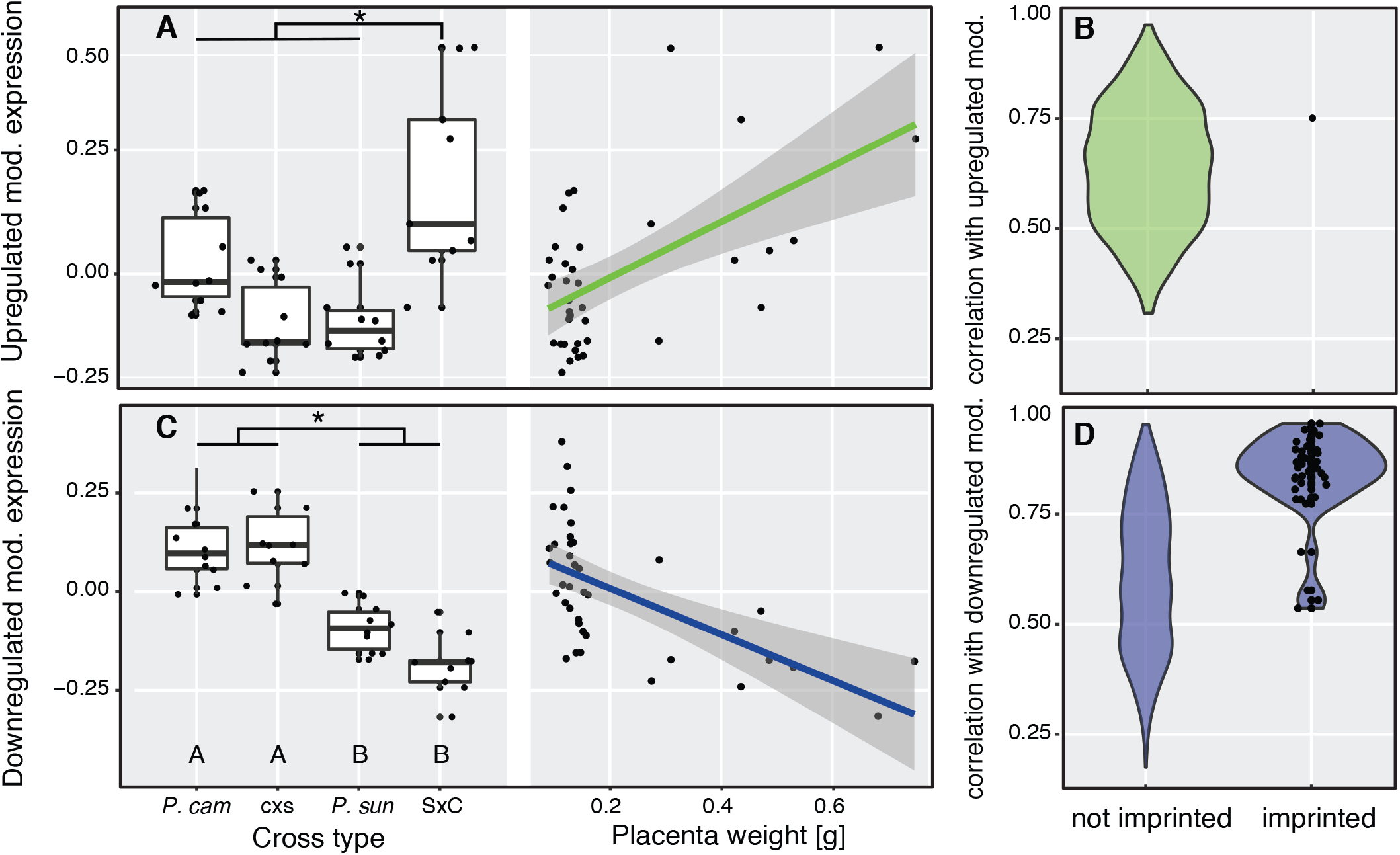
F_1_ placenta size-associated gene expression modules. Eigengene gene expression values summarize the group of genes that are upregulated (A, B) and downregulated (C,D) in overgrown S×C placentas, shown as differing by cross type (transgressive expression; stars indicate significance groups at P < 0.05 based on a Tukey’s HSD test) and placental size (association with hybrid incompatibility phenotype controlling for Theiler stage and sex, P < 0.001, ANOVA, see Table S2). The upregulated F_1_module was not enriched for imprinted genes (C; total genes, N = 565; imprinted genes, N = 1), whereas the downregulated F_1_ module was enriched for highly connected candidate imprinted genes (Brekke *et al*. 2016) showing maternally biased expression (D; total genes, N = 1160; imprinted genes, N = 36). Data points rendered only for imprinted genes in B and D.

**Figure 4.**
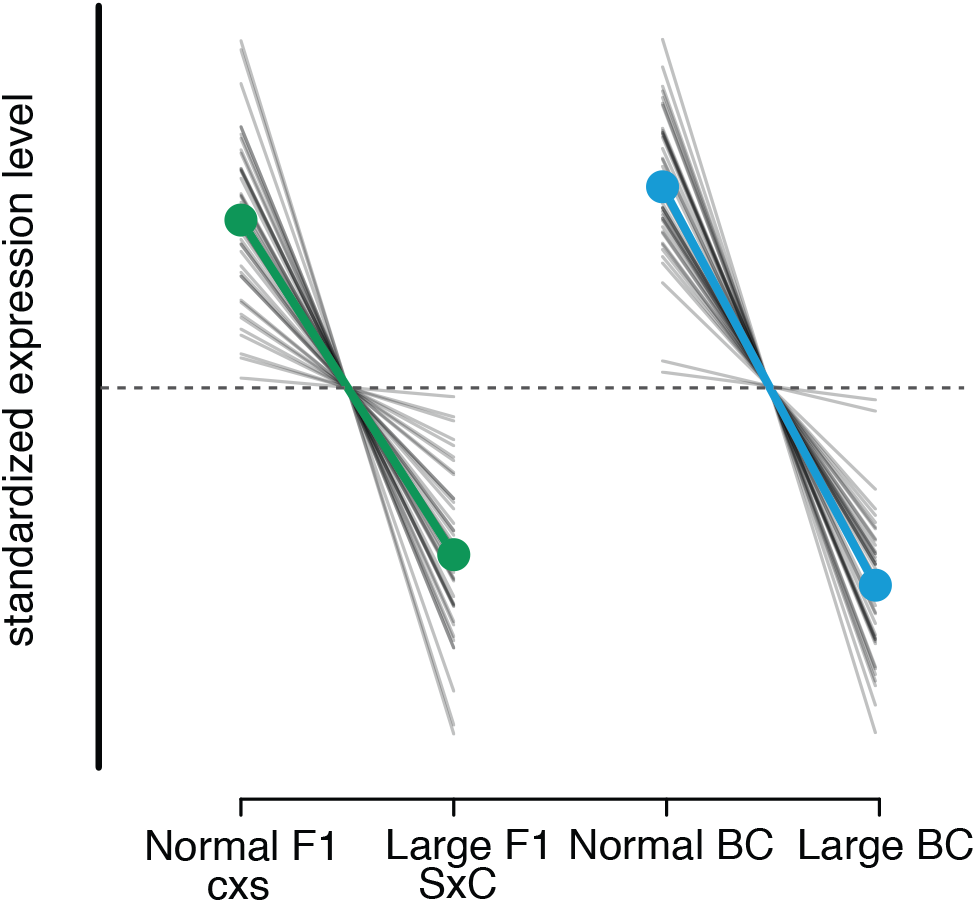
Reduction in candidate imprinted gene expression in overgrown F_1_ and BC placentas. Each gray line represents the relative change in expression of a candidate imprinted gene between normal sized and large hybrid placentas. Genes were standardized by mean expression per gene (dotted line), and centered to display direction rather than magnitude of gene expression change. Both F_1_ and BC experiments demonstrate reduced gene expression of genes showing parent-of-origin bias accompanying placental overgrowth (grand mean indicated by colored line).

### Networks of placental gene expression in second generation backcrossed hybrids

To test for links between our backcross QTL mapping experiment and emergent patterns of placental expression in F_1_ hybrid models, we next analyzed 24 transcriptomes from backcross placentas (12 large and 12 normal sized placentas; 11,396 genes placed in the network). One large female placenta was removed during outlier filtering (Figure S10). WGCNA analysis of the backcross transcriptomes revealed seven modules that were correlated with placenta weights. No clusters were significantly associated with embryo weights after controlling for developmental stage and sex (Table S3). The recombinant genotypes within this backcross sample allow us to more clearly differentiate disrupted expression in overgrown hybrid placenta versus species differences (*P. sungorus* versus *P. campbelli*) or interspecific hybridization per se (hybrids versus parental species). Consistent with this, only about one-third of the core genes included in the upregulated and downregulated F_1_ modules were captured in the seven placenta-associated backcross modules (563 of 1725 genes). However, most of these overlapping genes (>90%, 513 of 563) were derived from the downregulated F_1_ S×C module (Table S3). The backcross module that was most strongly associated with placental weights (432 genes) tended to show lower summary expression in overgrown backcross placenta (Figure 5A). This downregulated backcross module was also highly enriched for autosomal imprinting (33 candidate imprinted genes, binomial exact test P << 0.0001) and for genes in the downregulated _F_1 S×C module (206 genes, binomial exact test P << 0.0001; Table S3, Figure 4, Figure S11). In concordance with the overlap between gene lists, genes that were positively correlated with the F1 downregulated module eigengene were also positively correlated with the BC downregulated module eigengene (Figure S12), indicating module conservation across first and second generation hybrids.

**Figure 5.**
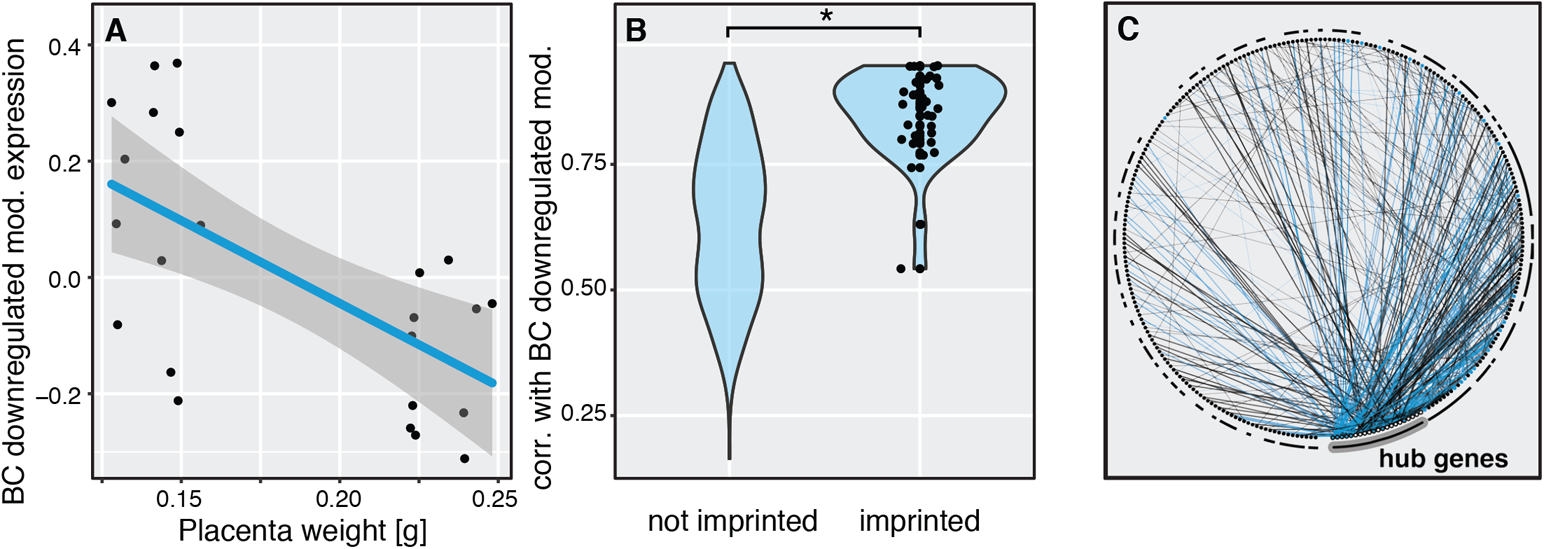
BC downregulated module is associated with placenta size and candidate imprinted genes. (A) Summary expression of the BC downregulated module was significantly associated with placenta size controlling for Theiler stage and sex (p < 0.001, ANOVA, see Table S3). (B) This module was enriched for highly connected, candidate imprinted genes (μ_imprinted_=0.8489±0.015; μ_not imprinted_=0.6370±0.008; Welch’s two-sided T-test, t_49.315_=12.255, P<0.0001; total genes, N = 432; imprinted genes, N = 33). Data points rendered only for imprinted genes. (C) The top 500 pairwise connections within this network largely involved the top 5% most connected genes, with interactions involving hub genes denoted by a thicker line and a solid black circumference notation at the interaction partner. Interactions involving a candidate imprinted gene are indicated in blue.

Connectivity within scale-free expression networks is commonly defined by the extent to which the expression level of a given gene is correlated with the expression of other genes. We found that candidate imprinted genes were much more highly connected than non-imprinted genes within the downregulated, placenta-associated backcross module (Figure 5B). Co-expression modules are usually characterized by a few highly connected “hub” genes (Ghazalpour *et al*. 2006; Mack *et al*. 2019). We found that the top 5% most connected (hub) genes in the downregulated module were involved in 60% (300) of the top 500 pairwise interactions (Figure 5C, hub interactions indicated by thicker lines and larger circles). Candidate imprinted genes were highly overrepresented as hub genes in this network — one third of hub genes (8 of 21 genes, binomial exact test P << 0.0001) were candidate imprinted loci with maternally-biased expression, including the top five most highly connected genes (*Plxdc2, ProcR*, Scara5, CD68, and *Wnt4*). Indeed, nearly half (238) of the top 500 pairwise correlations (Figure 5C, blue lines) involved at least one candidate imprinted gene (binomial exact test P << 0.0001). These highly-connected, downregulated genes represented many core biological functions of the placenta, ranging from broadly expressed genes involved in growth and development to those with specialized placental function (Table 1).

**Table 1.**
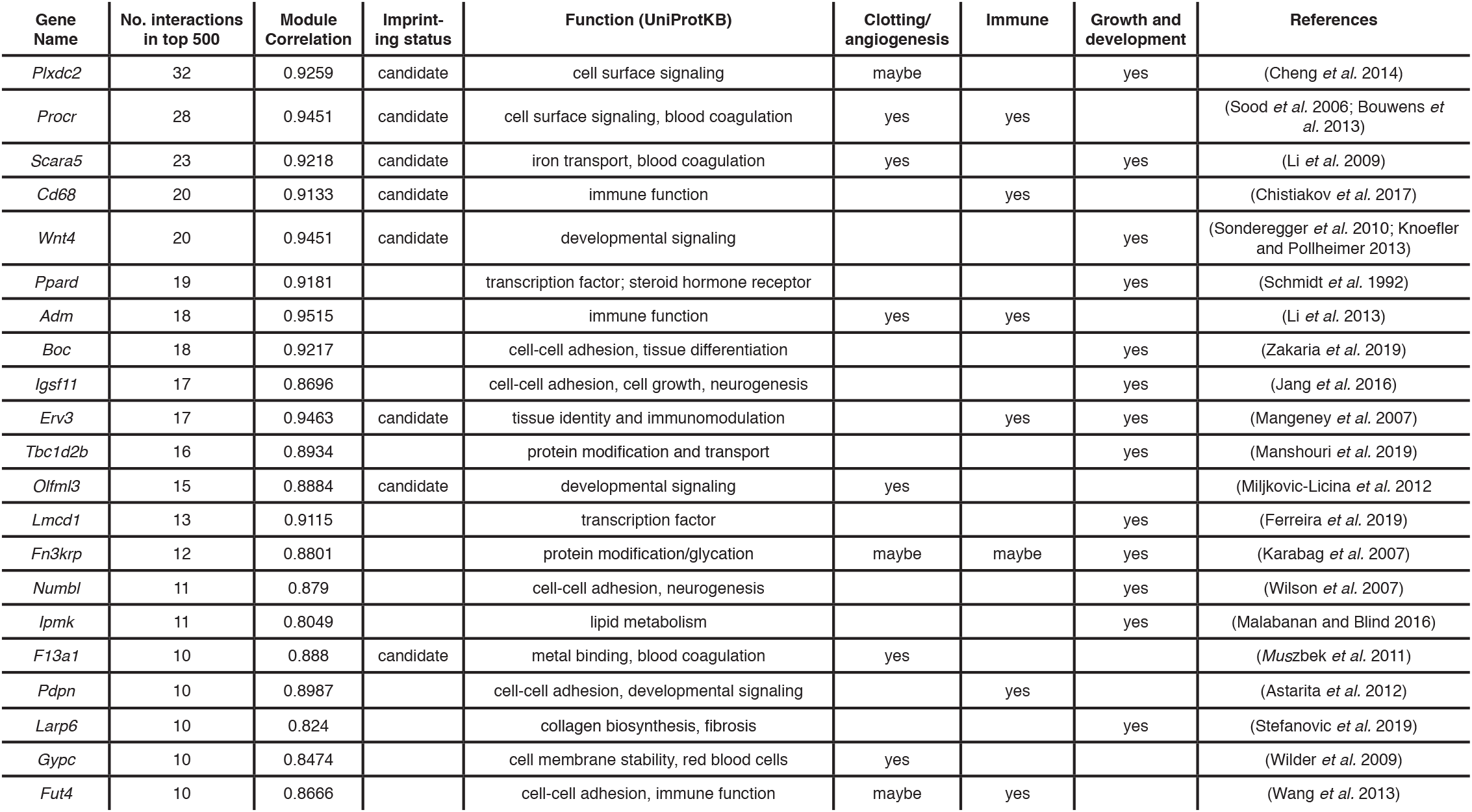
Hub genes in the downregulated, placenta-associated BC module. The top 5% most connected genes in the downregulated module, with connection defined as the number of times the gene was included in the top 500 strongest pairwise correlations in gene expression between genes in the module. Candidate imprinting status (Brekke *et al*. 2016) and function of the genes are indicated, with emphasis on placental functions of clotting and angiogenesis, immunity, and development.

Our transcriptome analyses revealed a central link between the X chromosome and the disruption of autosomal regulatory pathways in the placenta. To integrate our expression and placental phenotypes more directly, we next tested for QTL that explained expression variation in the overall module eigengene. Despite a small sample size (n=23), the X chromosome was a significant predictor of the downregulated BC module expression after permutation (Figure S13A). In principle, this signal could represent a predominant contribution of X-linked genes to the eigengene summary of expression within this parent-of-origin module, or a genome-wide trans-regulatory signal dependent on the species origin of the X chromosome. Consistent with the latter hypothesis, only ∼6% of genes in downregulated backcross module were X-linked (25 of 432 genes, binomial exact test P = 0.006, not significant after correction for multiple testing). These genes were significantly under-represented in the correlation network with only 7 of the top 500 pairwise correlations including an X-linked gene (56 pairs expected; binomial exact test P << 0.0001). It is also possible that the strong X-linked signal could be a correlated side effect of overall placental phenotypes on expression levels caused by X-linked hybrid incompatibilities. However, we found no significant QTL for the six other expression modules correlated with placental weight (Figure S13B,C), thus this signal seems unlikely to be a consequence of a spurious phenotypic correlation.

### Integration of the placental transcriptome onto the Phodopus genetic map

These results strongly support the hypothesis that X-linked incompatibilities interacting with a hybrid autosomal background are the primary determinant of disrupted autosomal expression observed in both F1 and BC hybrids, but the architecture of underlying X-autosome incompatibilities remain unresolved. The current genome assembly for dwarf hamsters is highly fragmented and has not been arranged into an ordered physical map (Bao *et al*. 2019), limiting our ability to fully integrate our transcriptome and quantitative genetic analyses. To begin to overcome these limitations, we used a custom exon capture experiment to anchor 3,616 placenta-expressed genes onto the *Phodopus* genetic map, including 159 X-linked and 34 autosomal imprinted genes (Table S13, Figure S14). An additional 212 X-linked genes were identified based on patterns of inheritance and orthology with mouse, but were not ordered on the genetic map (371 X-linked genes total). We placed approximately one-third of the BC downregulated network genes (162 of 432 genes) on to the genetic map, including 17 of the 21 hub genes (Table 1). Genes in this anchored network were distributed across 12 of the 13 autosomes and the X chromosome (Figure S15).

### A scan for genetic interactions that influence gene expression

If there was a hybrid interaction between the X chromosome and a specific diploid genotype at an autosomal locus that influenced the expression of that gene, then individuals that inherited a maternal *P. sungorus* X should show a larger change in gene expression for one of the autosomal genotypes than for the other. To test for such interactions, we calculated expression interaction (EI) scores by comparing the fold change difference in expression between the two possible autosomal genotypes of all mapped genes (i.e., homozygous for the *P. campbelli* allele or heterozygous) dependent on the genotype of the maternal X chromosome (*P. campbelli* or *P. sungorus*). Mapped genes from the BC downregulated module (n=124) showed higher EI scores on average when compared to genes not placed in any WGCNA module (mean_downregulated_=0.229±0.008, mean_null_=0.165±0.006, F_1,870_ = 14.13, P =0.0002, ANOVA). Next, we used a linear model to assess the relationship between gene connectivity and EI score in the downregulated module. Hub genes disproportionately overlap with male sterility loci in hybrid mice (Morgan *et al*. 2020), suggesting that incompatibilities may be more common among highly connected genes. However, we found support for only a slight increase in EI score for genes that were the most connected to the module (EI ∼ module correlation, adjusted r^2^ = 0.0243, F_1,122_ = 4.063, P = 0.046; Figure 6A). Interestingly, much of the signal for increased EI scores appeared to be driven by candidate imprinted genes rather than hub gene status per se (EI ∼ candidate imprinting status, adjusted r^2^ = 0.059, F_1,122_ = 8.75, P = 0.0037; EI ∼ hub, F_1,122_ = 0.005, P = 0.942; Figure 6B, Figure S16), again underscoring the central role of genes with maternally biased expression within the downregulated module. One such candidate imprinted (non-hub) gene was *Tfpi2* (Figure 6C). Individuals with an interspecific mismatch between the maternally inherited X chromosome (*P. sungorus*) and maternal allele at *Tfpi2* (*P. campbelli*) showed more than a two-fold (2.26x) decrease in expression compared to individuals with matching maternal genotypes.

**Figure 6.**
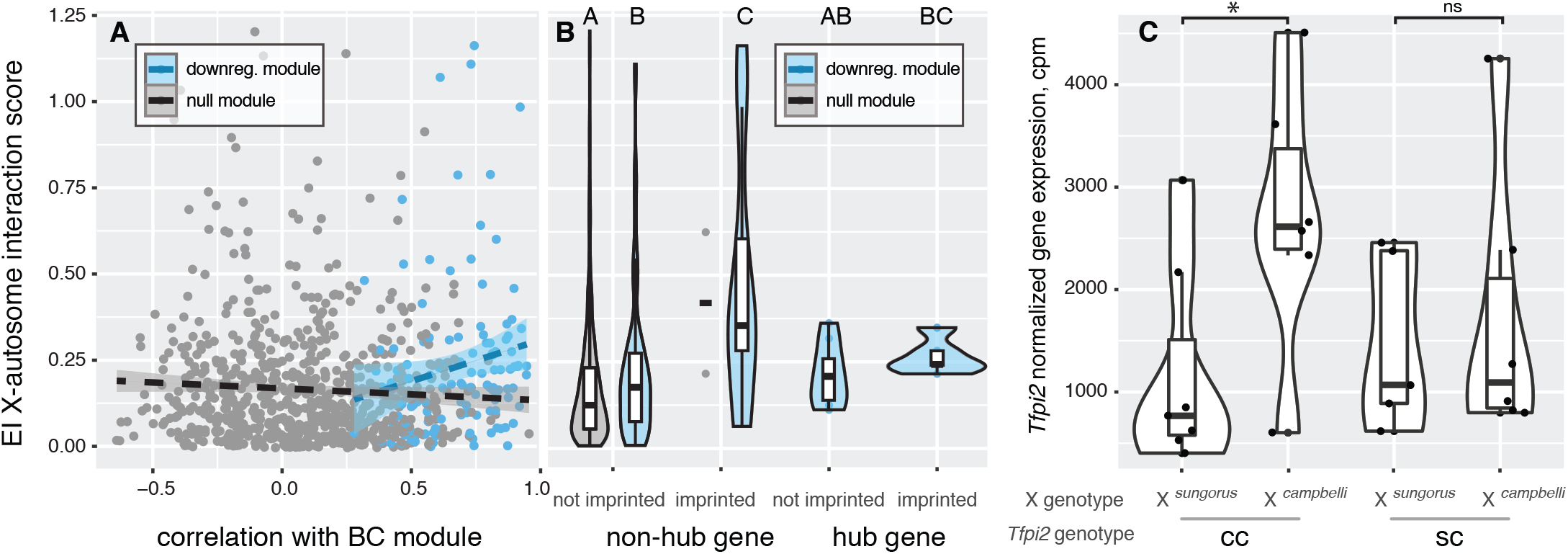
Expression interactions exposed in BC hybrids. (A) Greater disruption in gene expression with X chromosome – autosome mismatch (EI score) was weakly associated with genes more connected in the downregulated BC network module (F1,122 = 4.063, P = 0.046, ANOVA). Shown are comparisons between the gray module (gray; genes passing filter, N = 733) and the downregulated BC module (blue; genes passing filter, N = 124). The gray module includes all genes that were not placed in any module in the network analysis, and was used to generate a null expectation for the distribution of the EI score. (B) Increased EI score was explained by candidate imprinted genes, rather than highly connected hub genes (F1,122 = 8.75, P = 0.0037, ANOVA; downregulated BC module not imprinted non-hub genes, N = 102; imprinted non-hub genes, N = 9; not imprinted hub genes, N = 8; imprinted hub genes, N = 5). (C) EI expression pattern is illustrated with top gene Tfpi, where mismatch between maternal X genotype and maternal Tfpi genotype results in an average reduction in gene expression in addition to inheritance of a maternal *P. sungorus* X chromosome. Gene expression is reported as counts per million (CPM) transcripts. Letters and star indicate significant differences at P < 0.01, assigned by a Tukey’s HSD test.

We next polarized EI scores to evaluate specific X-autosome combinations with the downregulated module for candidate hybrid incompatibilities. Within the BC architecture, hybrid incompatibilities involving autosomal recessive or imprinted genes could manifest when the maternally inherited *P. sungorus* X chromosome was combined with maternally inherited *P. campbelli* autosomal alleles (all paternal alleles were *P. campbelli*). Consistent with this prediction, we observed positive polarized EI scores for imprinted genes within the downregulated module demonstrating that these expression interactions were driven largely by mismatches between the maternal autosomal and X chromosome genotypes (P = 0.043, Tukey’s HSD test, non-hub genes, imprinted vs null module; Figure S17A). Furthermore, we found that maternally mismatched imprinted genes in the downregulated module showed larger fold changes when compared to non-imprinted genes regardless of their status as a hub gene (P = 0.045, hub imprinted vs non-imprinted; P < 0.0001, non-hub imprinted vs non-imprinted, Tukey’s HSD test; Figure S17B). Collectively, these patterns suggest that imprinted autosomal genes with biased maternal expression within the downregulated module are more likely to be involved in incompatible hybrid interactions with the maternal X chromosome.

Finally, we tested for parent-of-origin bias in expression for candidate imprinted genes in the backcross. We found 31 genes previously identified as showing either maternal or paternal bias in expression in F_1_ placentas (Brekke *et al*. 2016) with expression data for at least seven heterozygous individuals in the BC. Of these, 22 genes show significant parent-of-origin bias in expression in the direction shown previously (Figure S18, one-sample T-test, Bonferroni corrected P < 0.05). Although limited to a subset of BC individuals, these results suggest that imprinting status was at least partially maintained at many genes despite large changes in overall expression levels. We also confirmed a strong bias in expression of the maternal allele on the X chromosome (adjusted r^2^ = 0.17, F_1,3560_ = 930.3, P << 0.0001; Figure S19), indicating that paternal imprinted XCI is likely intact in BC females as has been previously reported for F1 females (Brekke *et al*. 2016).

## Discussion

By combining quantitative genetic mapping of placental overgrowth with transcriptomic data, we uncovered genome-wide networks of gene expression that were disrupted as a consequence of incompatible genetic interactions with the X chromosome. These data indicate that genetic interactions between the X chromosome and networks of imprinted and non-imprinted autosomal genes are critical for proper placental development in dwarf hamsters. Below we compare our results with findings from other systems to examine the contribution of the X chromosome to the accumulation of reproductive barriers between species and the evolution of placental gene expression.

### The evolution of hybrid placental overgrowth

A positive parent-of-origin correlation between hybrid placental and embryonic growth effects has been observed in some mammal hybrids (e.g., Vrana *et al*. 2000; Brown *et al*. 2012), providing a possible mechanistic link between the disruption of early development and extreme adult parent-of-origin growth effects observed in many mammal hybrids (Brekke and Good 2014). Our quantitative results underscore that the relationship between placental and embryonic growth may be weak early in development, especially given the difficulty of differentiating the effects of edema and other developmental defects from overall embryonic growth (Figures S6 and S7). Consistent with this, parent-of-origin placental growth effects described in some mouse (Zechner *et al*. 1996; Kurz *et al*. 1999) and equine hybrids (Allen *et al*. 1993) also were not strongly correlated with hybrid embryonic growth.

Regardless of the unclear functional relationship between gross placental and embryonic growth phenotypes, our results show that the species origin of the maternal X chromosome is the major genetic factor responsible for placental overgrowth in *Phodopus* hybrids. Reproductive barriers between species often evolve through negative epistatic interactions between two or more loci, generally referred to Dobzhansky-Muller incompatibilities (DMIs; Dobzhansky 1937; Muller 1942; Orr and Turelli 2001, see also Bateson 1909). DMIs disproportionately involve X-linked loci, a phenomenon known as the large X-effect (Coyne and Orr 1989b). Hybrid incompatibilities presumably occur between the maternally inherited *P. sungorus* X chromosome and *P. campbelli* autosomal loci (e.g., dominant, recessive, or imprinted incompatibilities). However, paternal loci were invariant in our backcross and thus paternal contributions to putative X-autosome mismatches could not be mapped in this experiment. Large-X effects for disrupted placental development are perhaps not surprising given the central role that the X chromosome tends to play in the evolution of reproductive isolation (Coyne and Orr 1989b; Turelli and Orr 2000; Masly and Presgraves 2007; Turelli and Moyle 2007) and the parent-of-origin dependent nature of placental hybrid inviability in dwarf hamsters (Figure 2). However, we had previously associated placental overgrowth with widespread disruption of autosomal gene expression (Brekke *et al*. 2016); a regulatory phenotype that could manifest entirely from hybrid incompatible interactions between imprinted autosomal genes. Our mapping results rule out this possibility.

Large effect X-linked QTL also underlie placental overgrowth in hybrid deer mice (*Peromyscus maniculatus* x *P. polionotus*; Vrana *et al*. 2000) and house mice (*Mus* spretus x *M. musculus*; Zechner *et al*. 1996; Hemberger *et al*. 1999). In all three rodent systems, imprinted XCI occurs in the placenta and incompatibilities on the maternally inherited X chromosome emerge as a central genetic determinant of placental overgrowth in hybrids. However, broader connections between these developmental phenotypes and the disruption of placental gene expression have remained unclear. Studies in deer mice have linked the X chromosome with disrupted imprinted placental pathways, including a putative interaction with the autosomal maternally imprinted gene *Peg3* (Loschiavo *et al*. 2007). X-linked incompatibilities are also the primary cause of hybrid placental overgrowth growth in some hybrid crosses of house mice (*M. spretus* x *M. musculus*; Zechner *et al*. 1996; Hemberger *et al*. 1999), but no direct link between disrupted expression of candidate imprinted genes and placental overgrowth has been established (Shi *et al*. 2004; Zechner *et al*. 2004). However, a recent genome-wide study on the same *Mus* hybrid system showed transgressive autosomal expression in undergrown hybrid placentas (reciprocal F1 crosses were not performed), including disruption of the imprinted *Kcnq1* cluster (Arévalo *et al*. 2021). In these experiments, males showed more severe placental undergrowth and disrupted gene expression, which the authors proposed may involve interactions with the imprinted X chromosome (Arévalo *et al*. 2021). Artificial insemination experiments between more divergent *Mus* species pairs (*M. musculus* x *M. caroli*) resulted in massively abnormal placenta showing local demethylation and overexpression of an X-linked retroelement (Brown *et al*. 2012). Similarly, loss of genomic imprinting has been correlated with overgrowth in fetuses derived from assisted reproduction between divergent cattle breeds (Chen *et al*. 2015; Chen *et al*. 2016). Most of these works have focused on F1 crosses or candidate gene approaches, limiting further insights into the role that X-autosome interactions play in the broader disruption regulatory pathways in hybrid placenta. Building on these previous studies, we show that X-linked hybrid incompatibilities underlie the disruption of placental growth, with widespread effects on the misexpression of imprinted autosomal pathways.

Gene expression plays a central role in organismal development and morphological evolution (King and Wilson 1975; Carroll 2008; Sears *et al*. 2015), but the overall importance of regulatory incompatibilities to species formation has remained unclear (Butlin *et al*. 2012; Guerrero *et al*. 2016). In mammals, progress has been made in linking disruption of specific epigenetic regulatory mechanisms on the X chromosome (e.g., meiotic sex chromosome inactivation) to the evolution of hybrid male sterility during the relatively early stages of speciation (Bhattacharyya *et al*. 2013; Campbell *et al*. 2013; Davis *et al*. 2015; Larson *et al*. 2017). However, it has remained unclear if other developmental pathways are also predisposed to disruption in animal hybrids (Coyne and Orr 2004; Brekke and Good 2014). We suggest that placental development may represent a second developmental hotspot for the evolution of postzygotic reproductive isolation through the widespread disruption of gene expression.

There are several interesting parallels between the evolution hybrid male sterility and abnormal hybrid placental development in mammals. The X chromosome appears to play a central role in the genetic basis of both hybrid male sterility (Storchová *et al*. 2004; Good *et al*. 2008; Davis *et al*. 2015) and placental overgrowth (Figure 2; Zechner *et al*. 1996; Hemberger *et al*. 1999; Vrana *et al*. 2000). Both forms of postzygotic isolation manifest during developmental stages where the X chromosome is subject to chromosome-wide epigenetic silencing. Whereas meiotic sex chromosome inactivation may often be disrupted in the case of hybrid male sterility (Lifschytz and Lindsley 1972; Bhattacharyya *et al*. 2013; Larson *et al*. 2017), imprinted XCI appears to be maintained in hybrid hamster placentas (Figure S19; Brekke *et al*. 2016). Although the specific mechanisms underlying abnormal placental development are yet to be determined, tissues with imprinted XCI may select for the evolution of co-adapted epistatic networks of gene expression that are subsequently prone to disruption in hybrids (see below). Finally, aspects of placental development and spermatogenesis are both thought to evolve rapidly in response to genomic conflict (Haig 2000; Larson *et al*. 2018), which may contribute to the rapid evolution of hybrid incompatibilities (Crespi and Nosil 2013). It is well-established that hybrid male sterility tends to evolve rapidly in animals (Coyne and Orr 1989a; Wu *et al*. 1996), while hybrid growth effects have been detected across a broad range of evolutionary divergence in mammals (Brekke and Good 2014). All three systems where X-linked placental incompatibilities have been found (*Phodopus, Mus, Peromyscus*) involve crosses between relatively closely-related species pairs (Brekke and Good 2014), suggesting that placental hybrid inviability can evolve rapidly and contribute to the early stages of reproductive isolation.

Through analysis of these parallel systems of hybrid sterility and inviability, a trend is emerging where sex chromosome evolution and genetic conflict within regulatory systems appears to fuel divergence within these key developmental processes (Crespi and Nosil 2013; Larson *et al*. 2018), ultimately leading to the formation of reproductive barriers between species. Striking parallels also exist in plants, where hybrid seed inviability also evolves rapidly (Garner *et al*. 2016; Coughlan and Matute 2020) and has been linked to the intensity of parental conflict (Coughlan *et al*. 2020) and the disruption of imprinted gene expression in the extra-embryonic endosperm (Wolff *et al*. 2015).

### The evolution of placental gene expression networks

In both rodents and humans, fetal-derived trophoblast cells shape the vasculature at the maternal-fetal interface, allowing for nutrient transport and immune modulation (Gris *et al*. 2019). Notably, nine of the most connected genes in the BC downregulated module had functions related to coagulation and/or angiogenesis. Endothelial protein C receptor (*ProcR*) is an important anti-coagulant receptor in the trophoblast coagulation cascade (Bouwens *et al*. 2013). Allelic variants that result in under-expression of *ProcR* are associated with fetal loss in humans (Cochery-Nouvellon *et al*. 2009), and there is some evidence that maternal and fetal *ProcR* genotypes can interact to either prevent or induce placenta-mediated adverse pregnancy outcomes (Sood *et al*. 2006). In a healthy rodent placenta, the coagulation initiating tissue factor is counterbalanced by anti-coagulation proteins produced in differentiated syncytiotrophoblast tissue (Sood *et al*. 2006). Development fails without the early expression of the coagulation cascade in the placenta (Isermann *et al*. 2003). However, low levels of anticoagulants later in development are associated with preeclampsia and pregnancy loss (Ebina *et al*. 2015). Several of the hub genes (Table 1) are known to contribute to differentiation of placental layers. For example, *Wnt* signaling is broadly important in placentation and embryonic development, and *Wnt4* specifically may be involved in signaling between the fetal and maternal placental layers (Sonderegger *et al*. 2010; Knoefler and Pollheimer 2013). Another such specialized hub gene, *Erv3*, is part of a family of genes co-opted from endogenous retroviruses and are involved in immunomodulation, fusion and differentiation of trophoblasts (Mangeney *et al*. 2007), and are increasingly recognized for their role in regulating placental gene expression (Pavlicev *et al*. 2015; Chuong 2018). Similarly, the Plexin domain containing 2 gene (*Plxdc2*) encodes an endothelial cell-surface transmembrane receptor (Cheng *et al*. 2014) that is often co-expressed with *Wnt* signaling genes (Miller *et al*. 2007). Other candidate hub genes play roles in cell-cell adhesion and differentiation (Wilson *et al*. 2007; Jang *et al*. 2016; Zakaria *et al*. 2019), immune function (Astarita *et al*. 2012; Li *et al*. 2013; Wang *et al*. 2013; Chistiakov *et al*. 2017), nutrient metabolism and delivery (Schmidt *et al*. 1992; Li *et al*. 2009; Malabanan and Blind 2016), and transcriptional regulation (Ferreira *et al*. 2019).

Overall, our expression data suggest a strong connection between placental overgrowth, the maternally expressed (*P. sungorus*) X chromosome, and the imprinted expression of autosomal genes. Our candidate imprinted gene set included several genes known to be maternally (e.g., *Igf2, Mest, Peg3*) or paternally imprinted (e.g., *Axl, H19, Tfpi2, Wt1*) in mice, as well as several novel candidates including most of the hub genes (Table 1). Confirmation that these candidates reflect the evolution of novel parent-of-origin epigenetic silencing in *Phodopus* (e.g., through DNA methylation or other mechanisms) awaits detailed functional validation beyond the scope of the current study. Others have argued that contamination of maternal blood or tissue may often bias patterns of allele-specific expression in the post-embryonic placenta (Wang *et al*. 2011, but see Finn *et al*. 2014). We previously found little evidence for extensive maternal contamination of dissected placental tissue in *Phodopus* (i.e., genome-wide paternal:maternal allele ratios were ∼1:1 in overgrown SxC placentas), but it is possible that maternally-biased expression of some of these candidates reflects a large maternal contribution to overall placental expression levels. Indeed, hub genes such as *Wnt4* are thought to be directly involved in signaling between the fetal and maternal placental layers (Sonderegger *et al*. 2010; Knoefler and Pollheimer 2013). Regardless of the underlying regulatory mechanisms – epigenomic imprinting or fetal-maternal transcript sharing – our results suggest that X-linked and autosomal genes with maternally-biased expression play a central role in the evolution of placental development and the disruption of placental pathways in hybrids.

The existence of placental networks of maternally-biased gene expression is consistent with some predictions of the co-adaptation theory of gene expression, whereby maternal expression at one gene can select for maternally-biased expression at other positively interacting genes (Wolf and Hager 2006; Wolf 2013; Wolf and Brandvain 2014; O’Brien and Wolf 2017). Such a co-evolutionary process should result in the broader integration of imprinted gene networks, the evolution of separate co-expressed networks of maternally and paternally expressed genes, and the exposure of epistatic DMIs in second (or later) generation hybrids with recombinant genotypes (Wolf and Brandvain 2014; Patten *et al*. 2016). We uncovered evidence for many such interactions in our preliminary screen for genetic interactions that influence gene expression levels in backcross placenta (Figures 6, S16, S17). For example, the maternally expressed *Tfpi2* gene showed a greater than two-fold decrease in expression when a maternally inherited *P. campbelli* allele was combined with a maternally inherited *P. sungorus* X chromosome (Figure 6C). *Tfpi2* is imprinted in the placenta where its expression may limit trophoblast invasion (Jin *et al*. 2001) and also downregulated in several types of cancer (i.e., a potential tumor suppressor; Konduri *et al*. 2001; Ribarska *et al*. 2010). More data are needed to determine if this and other genes showing significant expression interactions contribute directly to abnormal placental growth phenotypes in hybrids.

More generally, we propose that the X chromosome is likely to play a central role in the evolution of maternally-biased placental networks in many eutherian mammals. The strength of this prediction is dependent on patterns of X inactivation in the placenta and other extra-embryonic tissues of females as males only have a maternally-derived X. The paternal X chromosome appears to be silenced in the placenta and other extra-embryonic tissues in at least four genera of rodents (i.e., *Mus*, Takagi and Sasaki 1975; *Rattus*, Wake *et al*. 1976; *Phodopus*, Brekke *et al*. 2016; *Peromyscus*, Vrana *et al*. 2000), resulting in predominantly maternal expression of X-linked genes in the placenta of males and females (Dupont and Gribnau 2013; Lee and Bartolomei 2013). Imprinted XCI is expected to contribute the vast majority of maternally expressed genes in the rodent placenta, which as a consequence should favor the evolution of maternal expression at interacting autosomal genes. In contrast, XCI appears to be random in extra-embryonic tissues of humans (Moreira De Mello *et al*. 2010), cattle (Chen *et al*. 2016), pigs (Zou *et al*. 2019), horses (Wang *et al*. 2012), and possibly rabbits (Okamoto *et al*. 2011). The more frequent occurrence of random XCI in this limited sample suggests that imprinted XCI may have evolved more recently in rodent extraembryonic tissues (Okamoto *et al*. 2011), and therefore may not apply broadly across the radiation of placental mammals. However, it is worth noting that rodents comprise over 40% of all placental mammal species (Burgin *et al*. 2018). The current sample of XCI in extraembryonic tissues is also biased towards a just few major lineages (i.e., ungulates, primates, rodents, lagomorphs) and thus insufficient for accurate reconstruction of the ancestral state of extra-embryonic XCI in placental mammals. Moreover, male hemizygosity in species with random XCI may still favor the evolution of maternal expression at interacting autosomal genes under some conditions (Wolf and Brandvain 2014). Additionally, the physiological integration of maternal blood supply and trophoblast-generated fetal vasculature is a particularly compelling biological context that could favor the evolution of coordinated maternal-fetal gene expression networks, regardless of the pattern of XCI. Given our data and these general theoretical predictions, the broader relevance of X-autosomal gene expression networks to placental evolution and development warrant further consideration.

## Acknowledgements

Ryan Bracewell, Zak Clare-Salzler, Ted Cosart, Kris Crandell, Doug Emlen, Mafalda Ferreira, Lila Fishman, Evgueny Kroll, Lindy Henry, Matt Jones, Sara Keeble, Erica Larson, John McCutcheon, Colin Prather, Brice Sarver, Vanessa Stewart, Dan Vanderpool, Jon Velotta, Paul Vrana, and the UNVEIL network provided helpful comments on data analysis and interpretation. Kelly Carrick, Jess Wexler, and the University of Montana LAR staff for helping with animal care. This research was supported by grants from the Eunice Kennedy Shriver National Institute of Child Health and Human Development of the National Institutes of Health (R01-HD073439, R01-HD094787 to JMG), the National Science Foundation (EPSCoR OIA-1736249 to JMG and ZAC), a National Science Foundation Postdoctoral Fellowship in Biology (DBI-1612283 to SCS), a National Science Foundation Doctoral Dissertation Improvement Grant (DEB-1406754 to TDB), a Society for the Study of Evolution Rosemary Grant Award (to TDB), and a David Nicholas Award (to TDB). This study includes research conducted in the University of Montana Genomics Core, supported by a grant from the M. J. Murdock Charitable Trust. The authors declare no competing financial interests.

---

**Figure S1.**
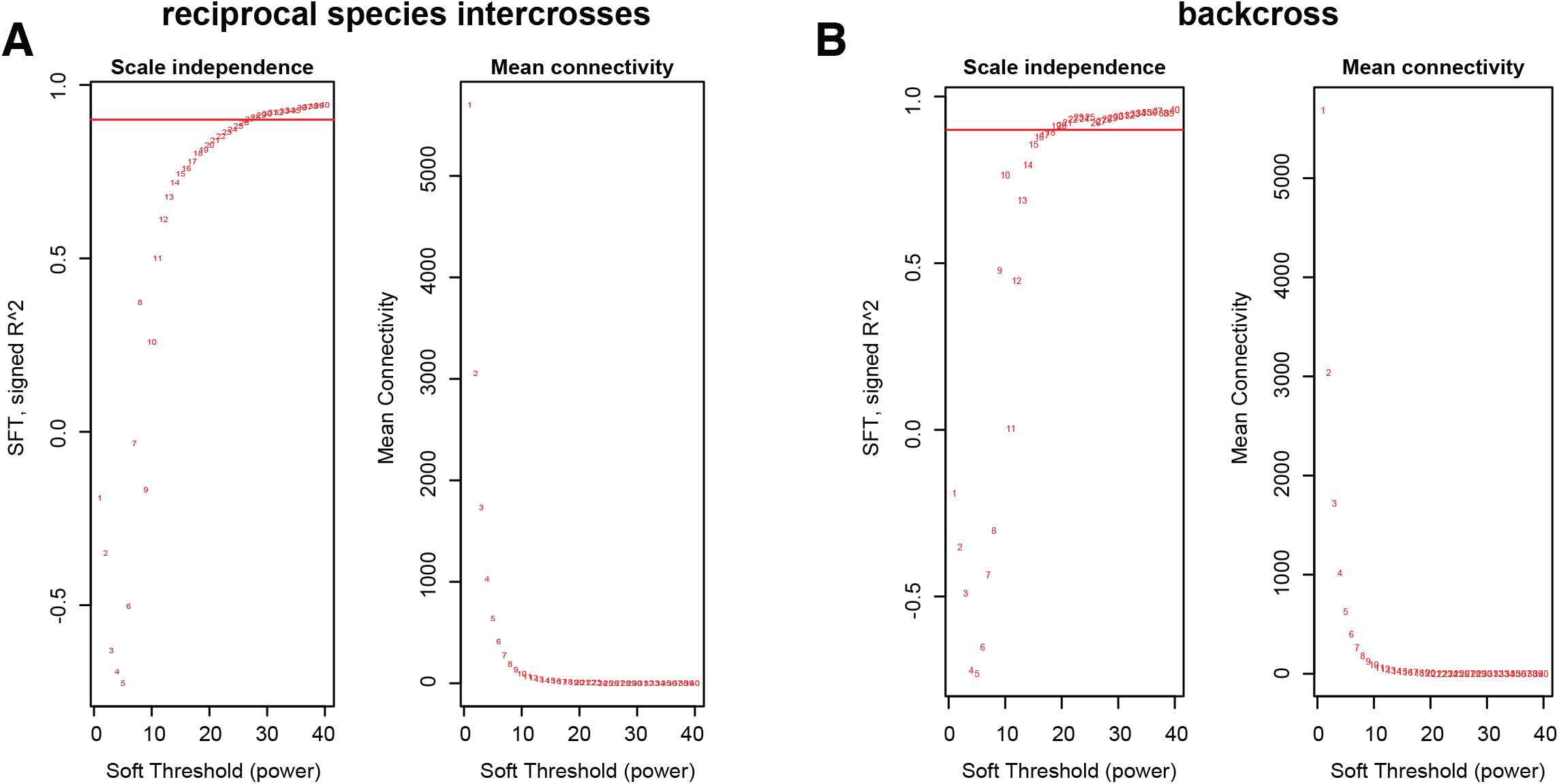
Soft thresholding power. Soft thresholding results for parental and reciprocal species intercross (A) and backcross (B) RNAseq datasets. These differ for each dataset, with the line indicating the threshold used in WGCNA cluster generation.

**Figure S2.**
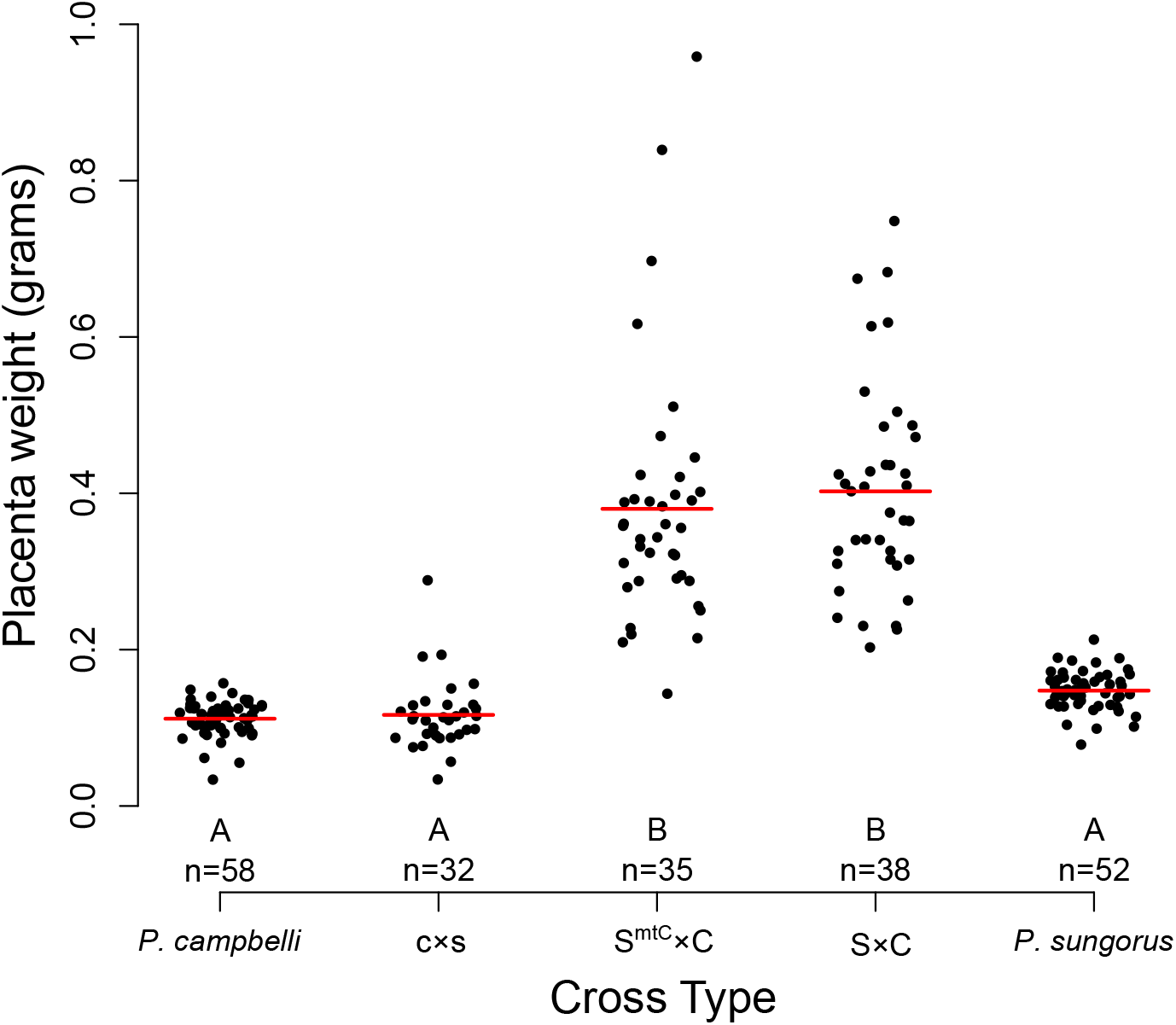
Mitochondrial interactions had no effect on placenta size. Placentas from the S^*mtC*^×C cross were indistinguishable from S×C hybrids (*F*_4,213_ = 106, P < 0.001, full ANOVA model, Tukey’s HSD test significance groups indicated by letter). Data for *P. campbelli*, c×s, S×C, and *P. sungorus* from (Brekke and Good 2014).

**Figure S3.**
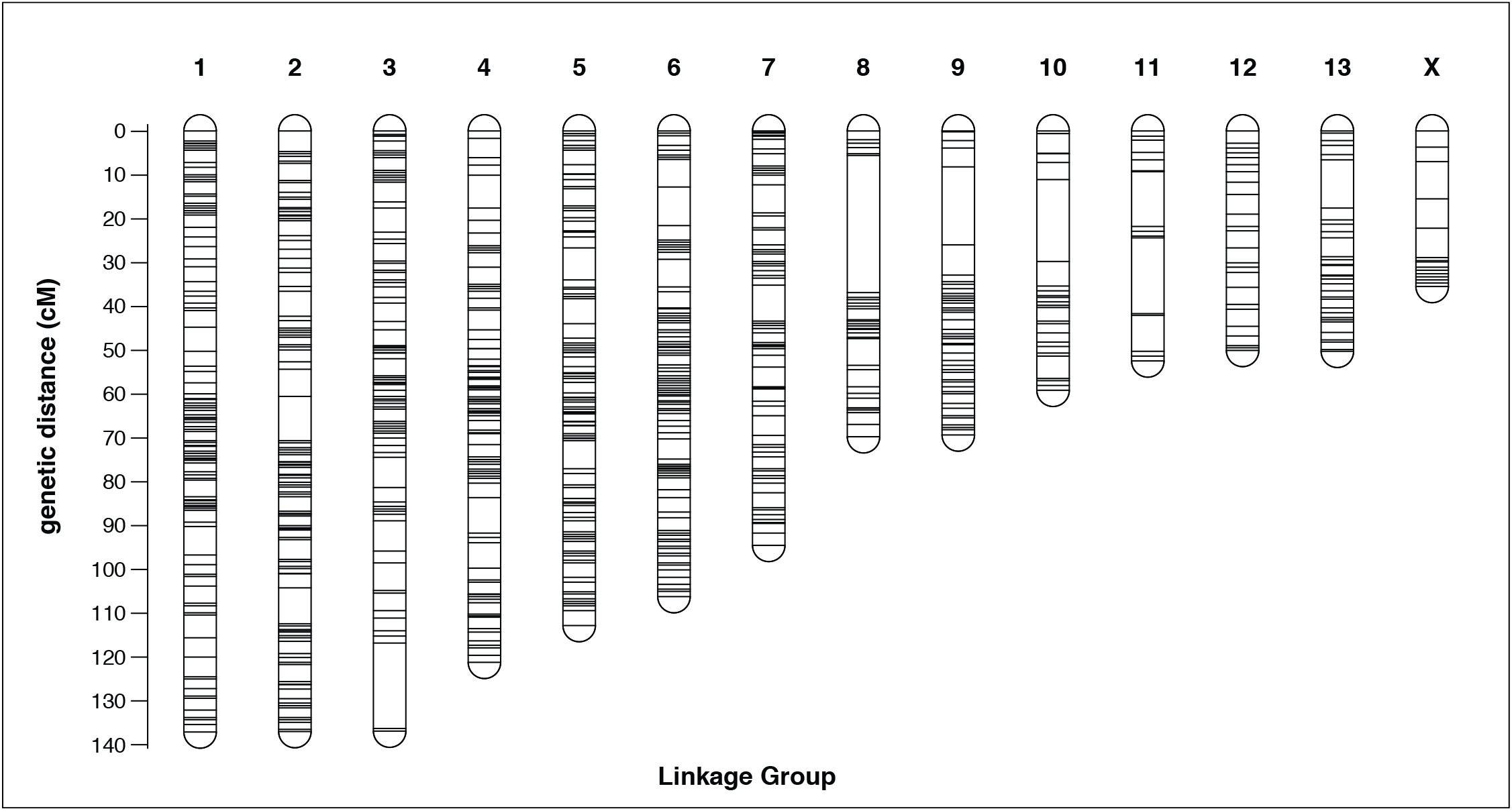
Genetic map of Phodopus dwarf hamsters. Map based on 1,215 RAD markers spanning 1,213.7 cM across 14 linkage groups numbered by decreasing length. The sequence of each marker and their exact locations in centiMorgans can be found in Supplemental Table 1. Specific locations of genes can be found in Table S2. Visualized with R/LinkageMapView (Ouellette et al. 2018).

**Figure S4.**
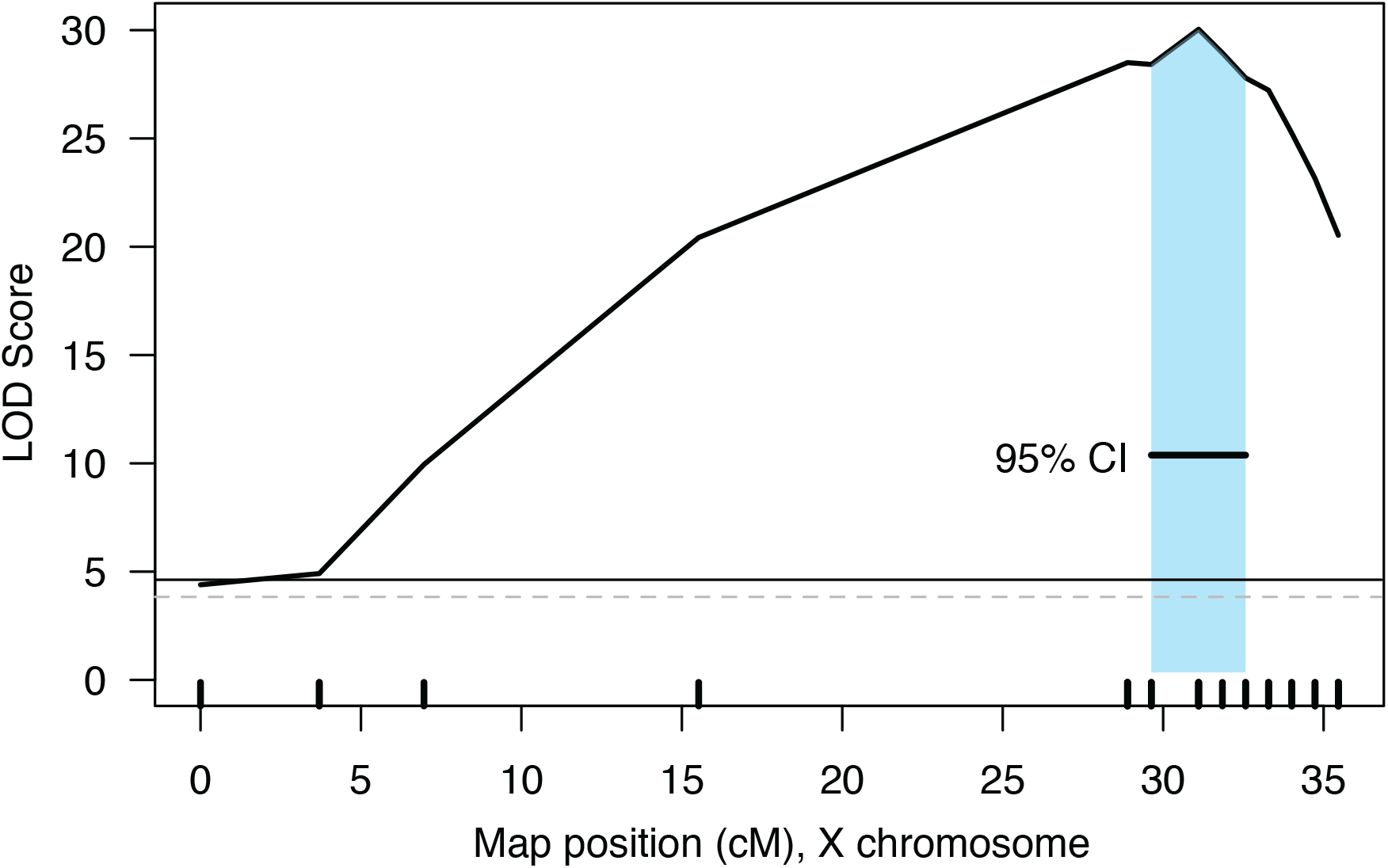
QTL interval on the X chromosome overlaps with increased marker density. Placental weight QTL likely corresponds to region of reduced recombination on the map. Solid line indicates permutation-based P = 0.01 significance threshold, dashed line indicates permutation-based P = 0.05 significance threshold.

**Figure S5.**
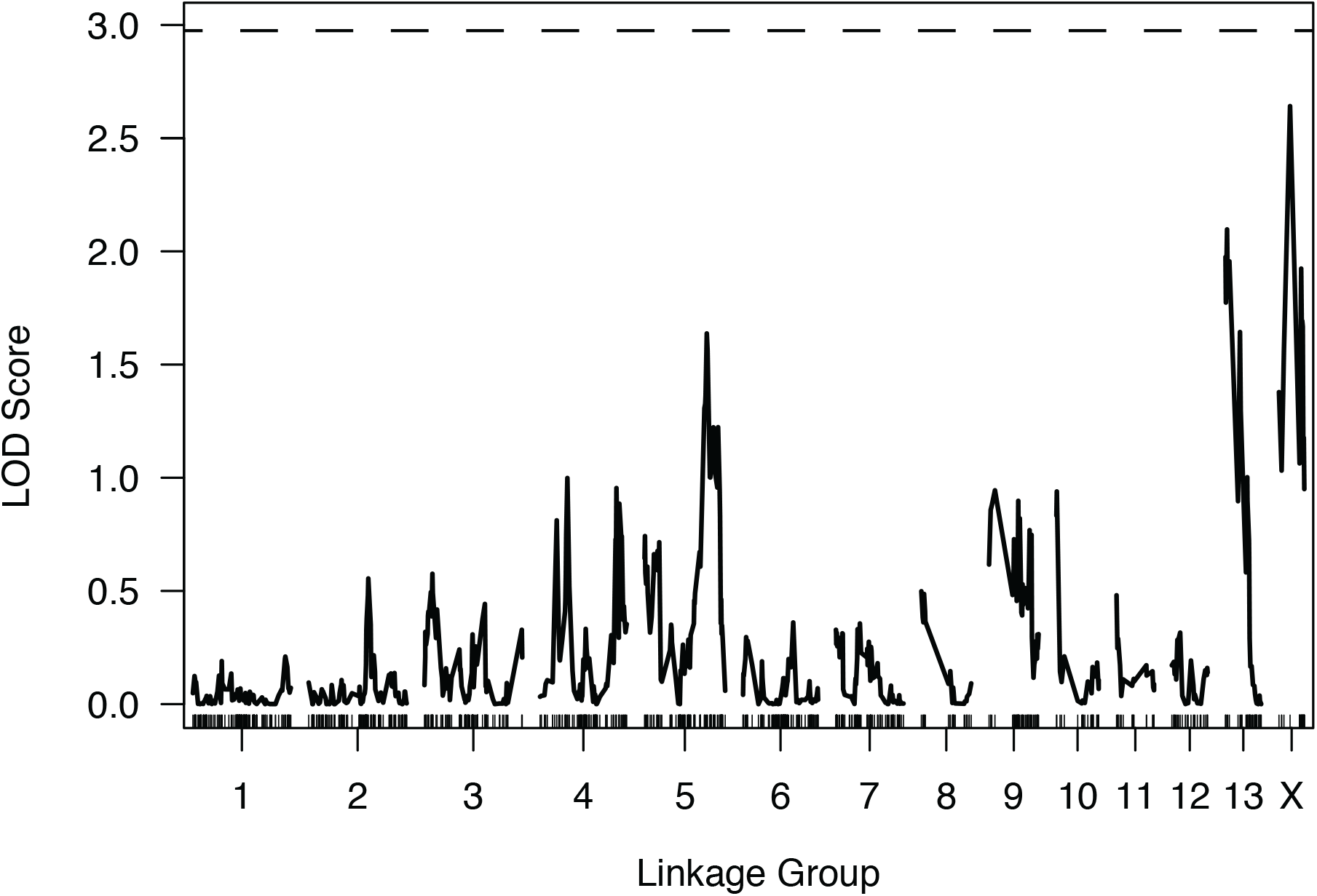
No QTL for embryo weight were detected in the BC mapping experiment. No peak passes the permutation threshold when controlling for Theiler stage and edema. P = 0.05 permutation threshold indicated with dashed line

**Figure S6.**
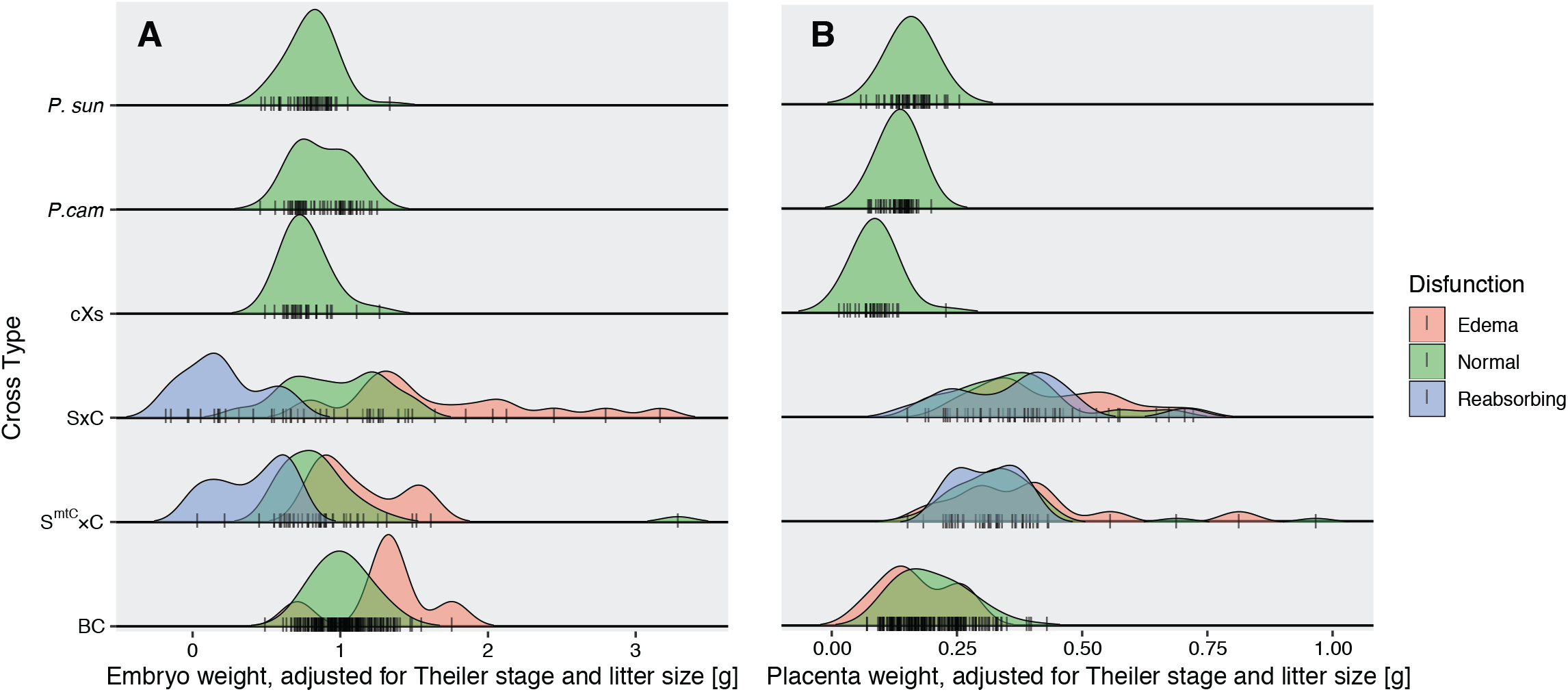
Edema and reabsorption explained much of the variance in embryo size, but not placenta size, in interspecies hybrid hamsters. Reabsorption (blue) and edema (red) shifted embryo size away from the mean in *P. sungorus x P. campbelli* (S×C) F, S^*mtC*^×C F1, and BC hybrid hamsters (A), but had little effect on placenta size (B).

**Figure S7.**
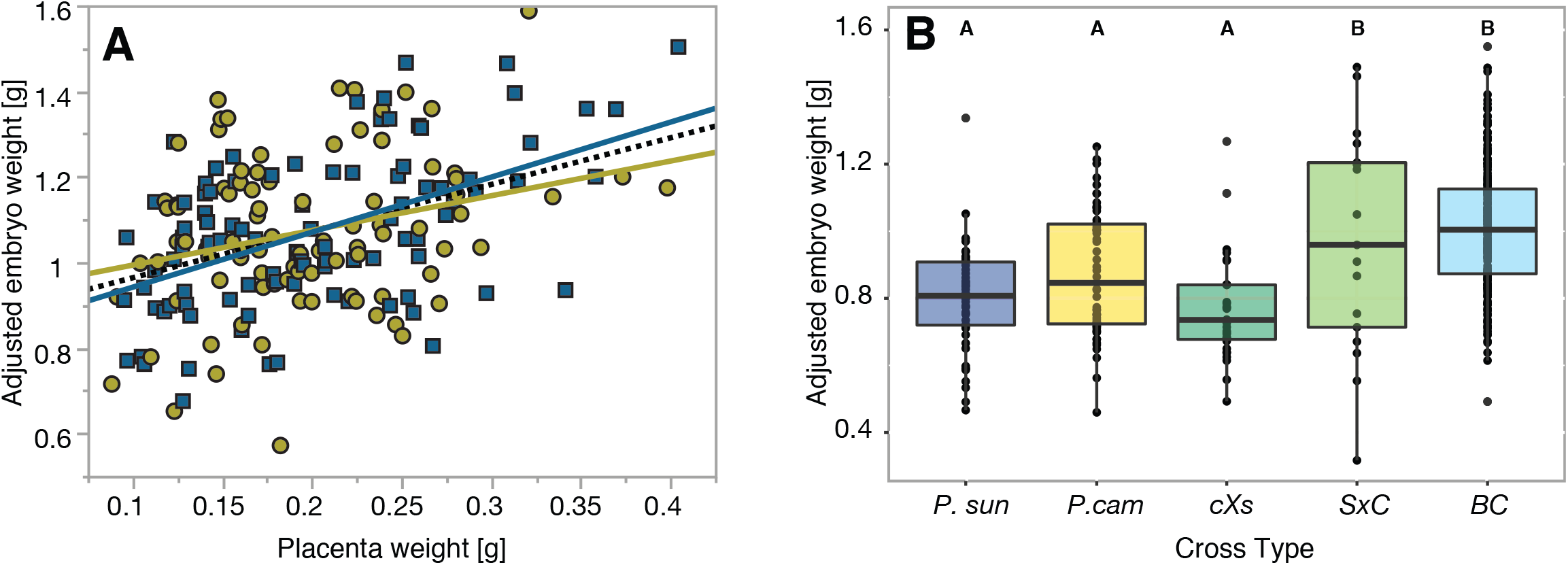
Embryo weight in the BC. (A) Embryo weight was positively associated with placental weight in the BC, and more strongly so in males (blue, adjusted r^2^ = 0.257, *F*_1,95_ = 33.8, P < 0.0001) than females (yellow, adjusted r^2^ = 0.065, *F*^1,88^ = 7.15, P = 0.0090). (B) When BC embryo weights were analyzed along with F1 hybrids, the overgrown SxC F_1_ and BC hybrids showed a slight but significant increase in size controlling for stage and edema (adjusted r^2^ = 0.159, *F*_1,184_ = 36.0, P < 0.0001, ANOVA).

**Figure S8.**
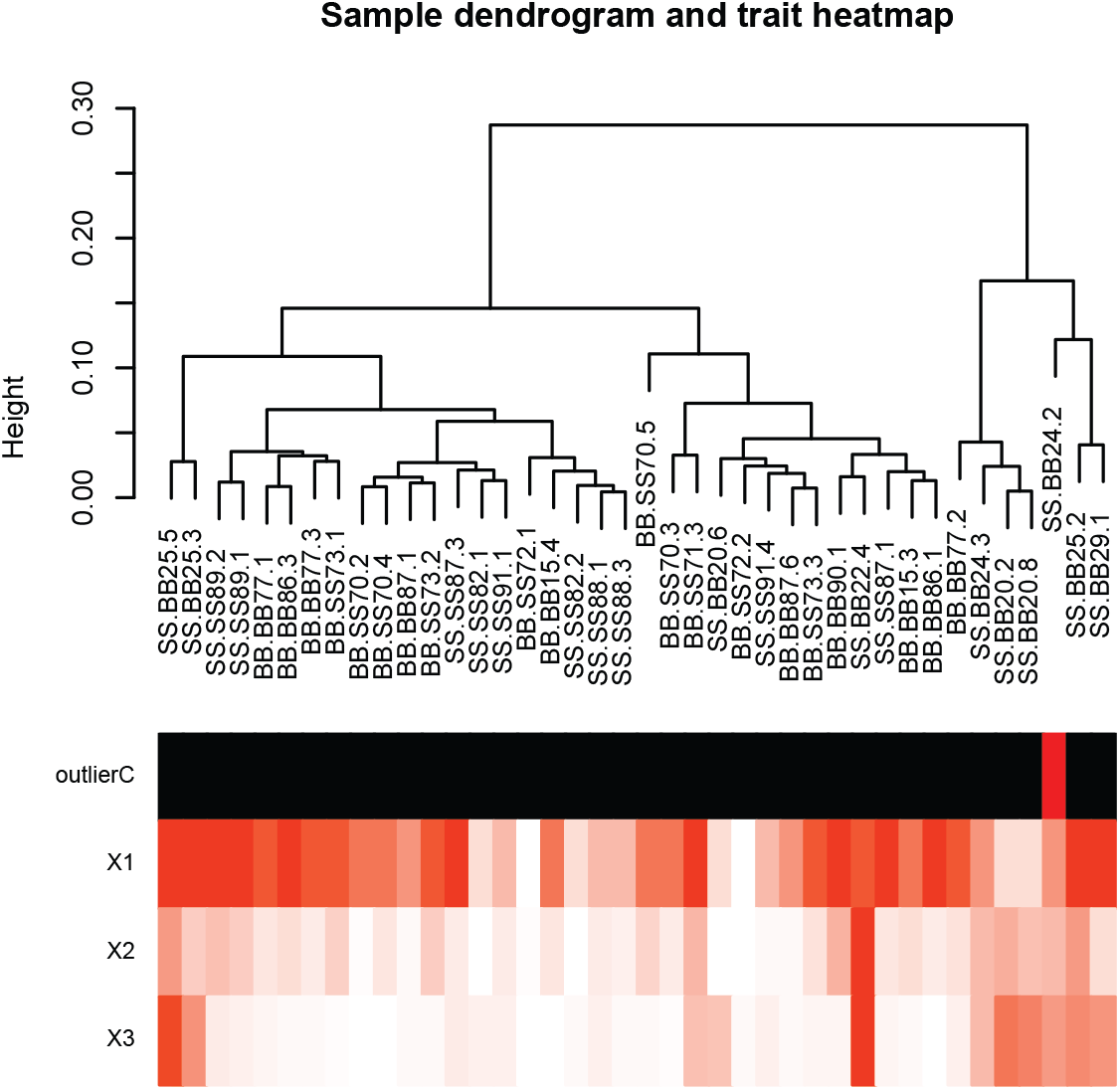
WGCNA output for F_1_ data. Outlier identification of sample to be removed in red.

**Figure S9.**
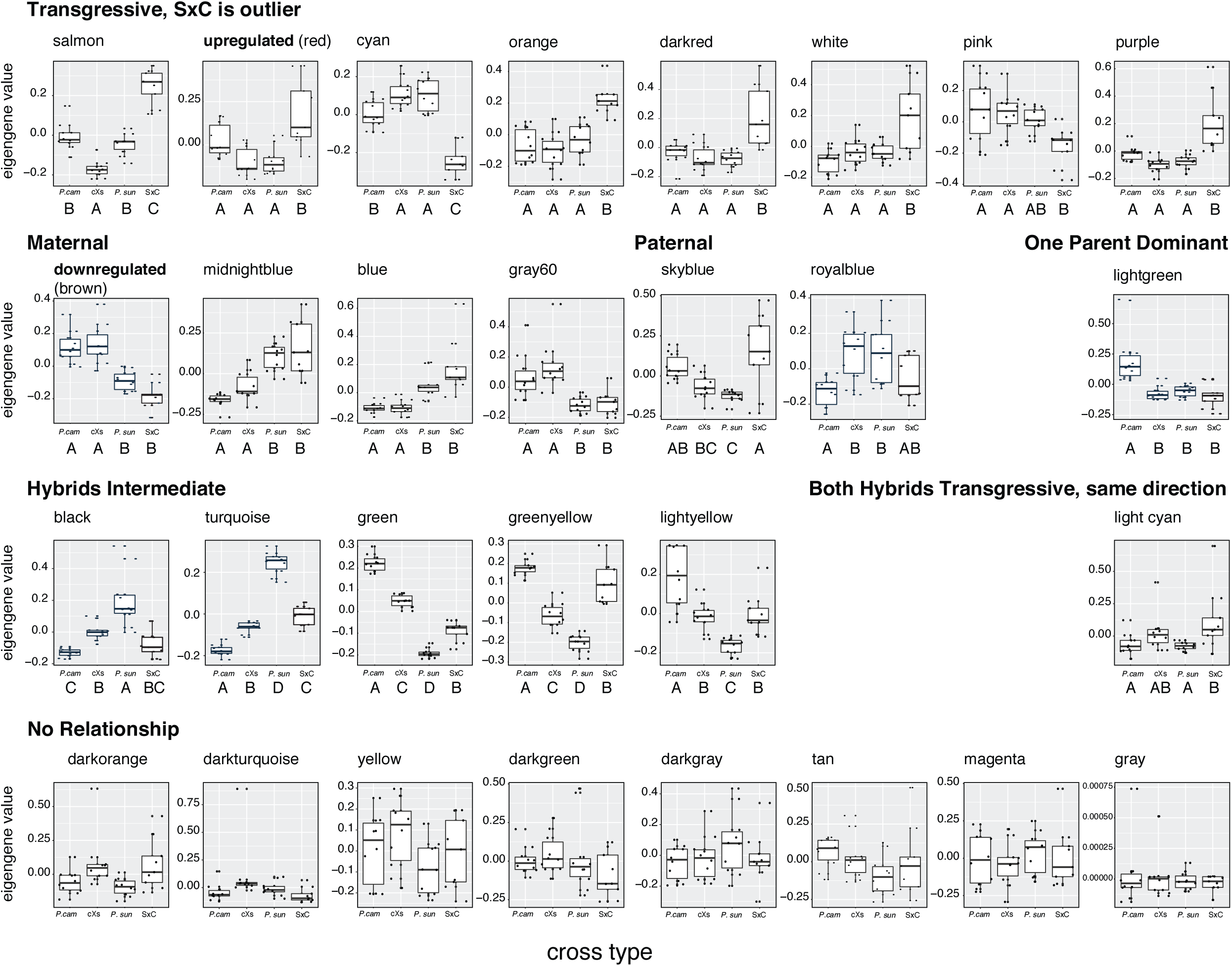
Assignment of inheritance patterns of modules in network. Summary values of module expression (module eigengene) tested with an ANOVA to identify parent of origin and transgressive expression. Color names are arbitrarily and randomly generated by the program, and have no additional meaning. Letters indicate Tukey’s HSD test assigned significance groups at P < 0.05.

**Figure S10.**
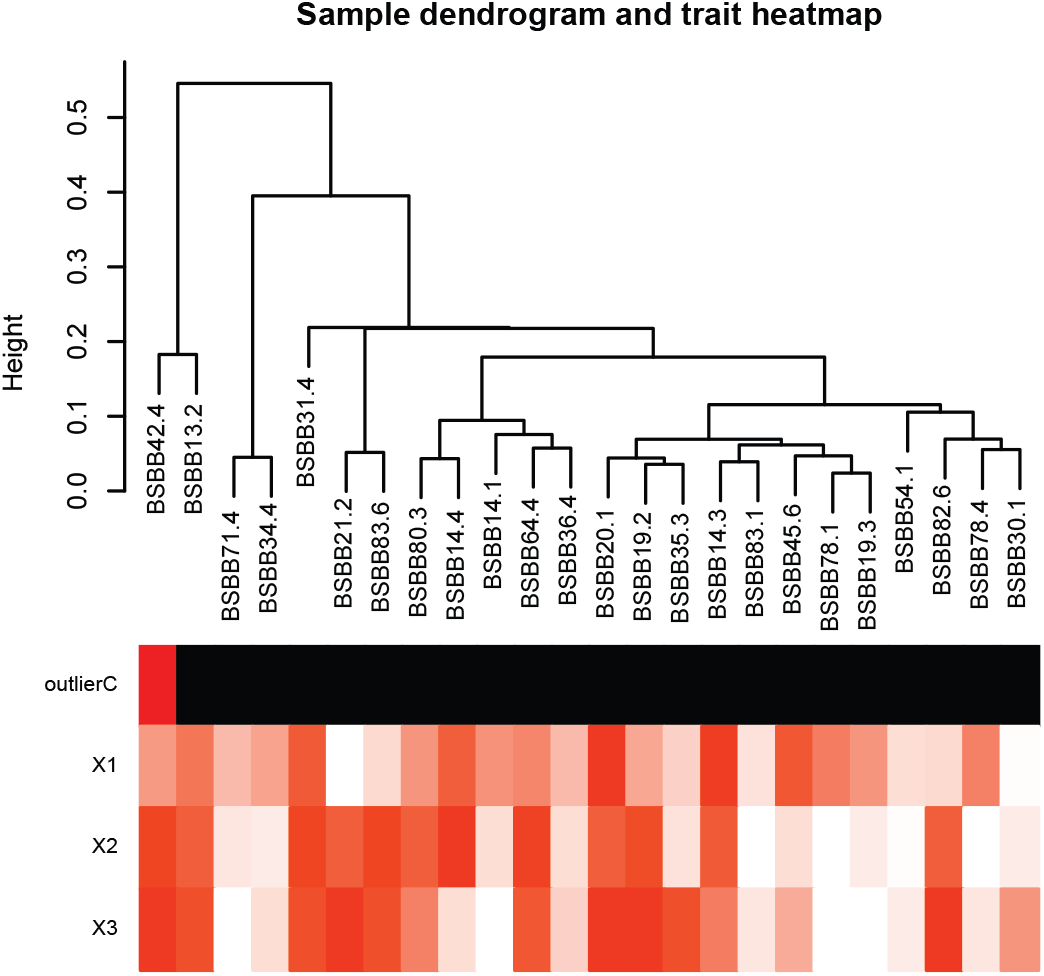
WGCNA output for BC data. Outlier identification of sample to be removed in red.

**Figure S11.**
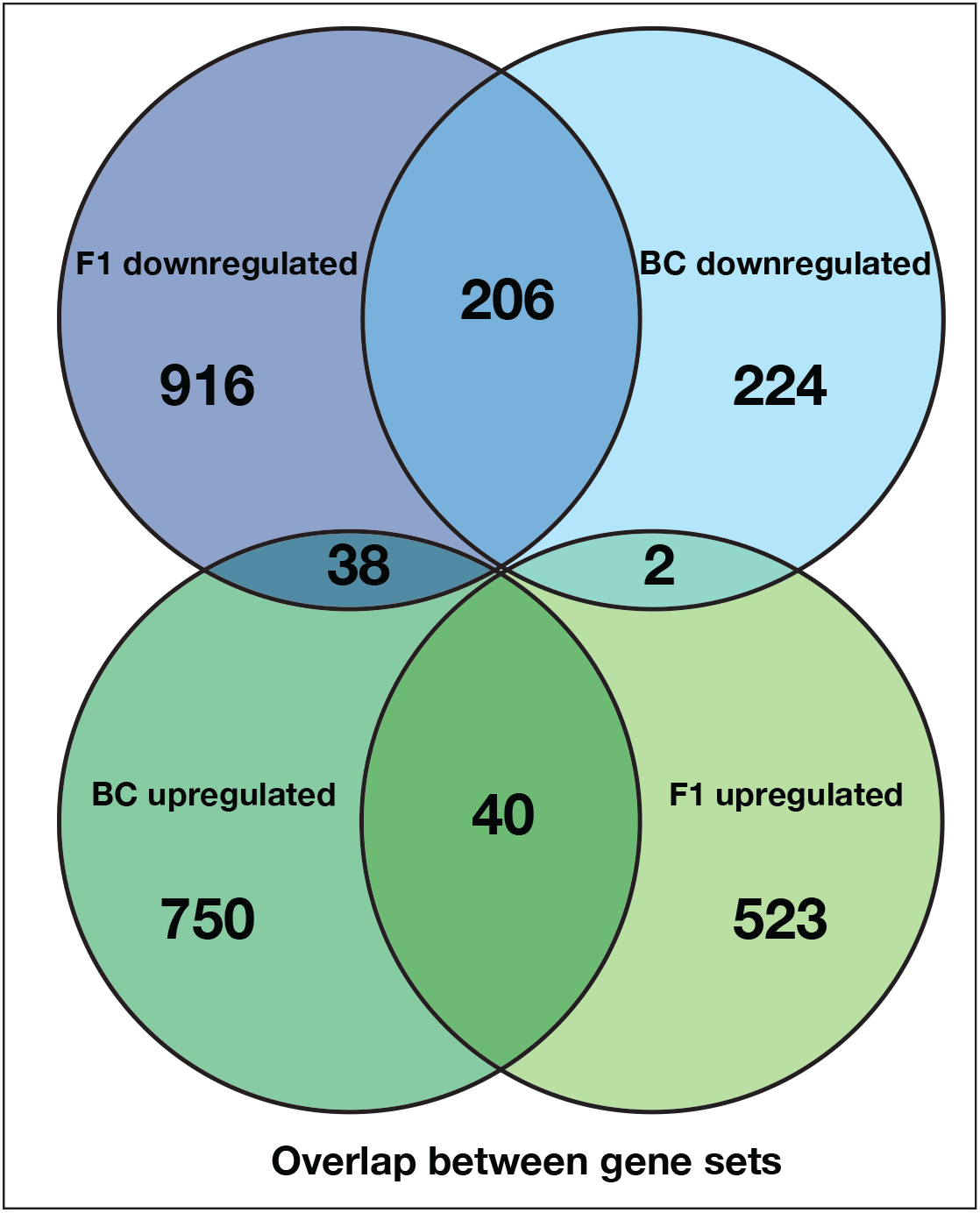
Count overlap between key F1 and BC modules. Venn diagram showing counts of genes shared between F_1_and BC downregulated and upregulated placenta modules. The downregulated modules shared the most genes (206).

**Figure S12.**
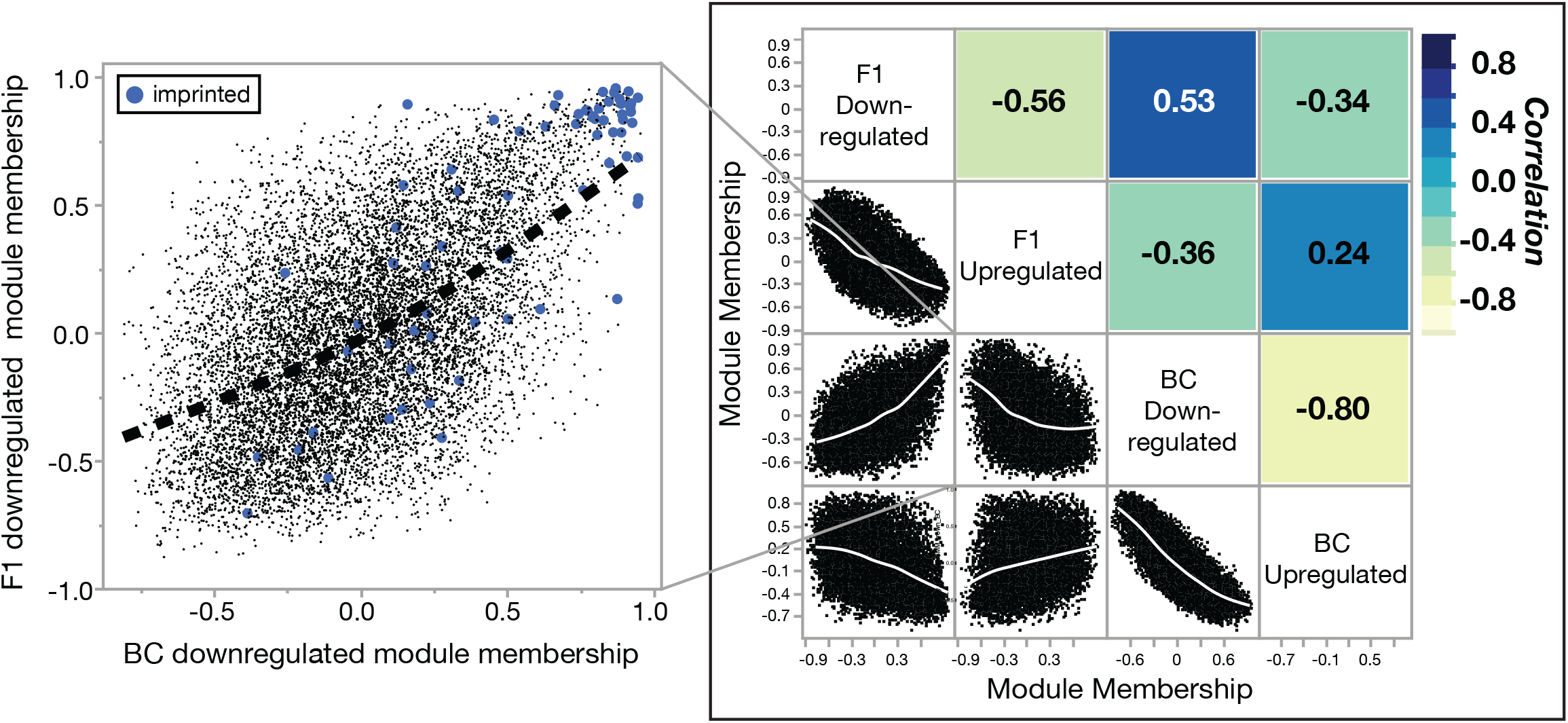
Correlation between key F1 and BC modules. Each gene in the network has a correlation to each module eigengene, whether that gene is placed in the module or not. We can assess how similar two modules are by asking whether genes are generally showing the same bivariate correlation to each module. Not only were the downregulated and upregulated modules within each data set negatively correlated with each other (F_1_, Pearson’s R = -0.56, P < 0.0001, BC, Pearson’s R = -0.80, P < 0.0001), the F_1_ and BC downregulated modules across experiments were positively correlated with each other (Pearson’s R = 0.53, P < 0.0001). Notably, the same candidate imprinted genes shared high connectivity/module membership with the network in both data sets (blue dots, inset).

**Figure S13.**
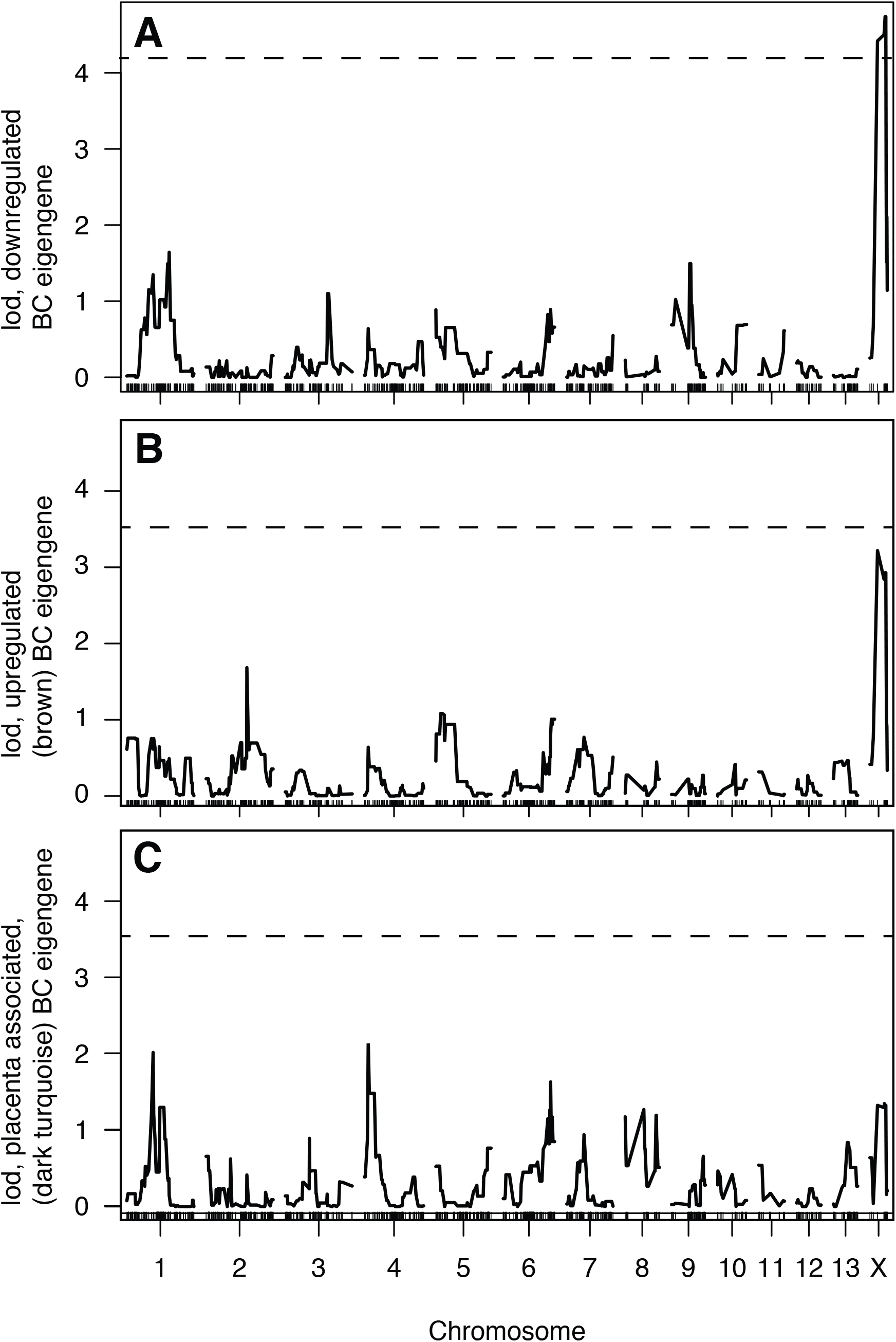
EigenQTL in BC hybrids. Using module eigengenes as phenotypes summarizing gene expression patterns in 23 BC hybrids, (A) we found that the downregulated module had a single X-linked QTL that passed a P = 0.05 permutation threshold (QTL peak at 31.1cM, LOD=4.739). (B,C) No QTL were detected for the other tested modules. Dashed line indicates permutation-based significance threshold (P = 0.05).

**Figure S14.**
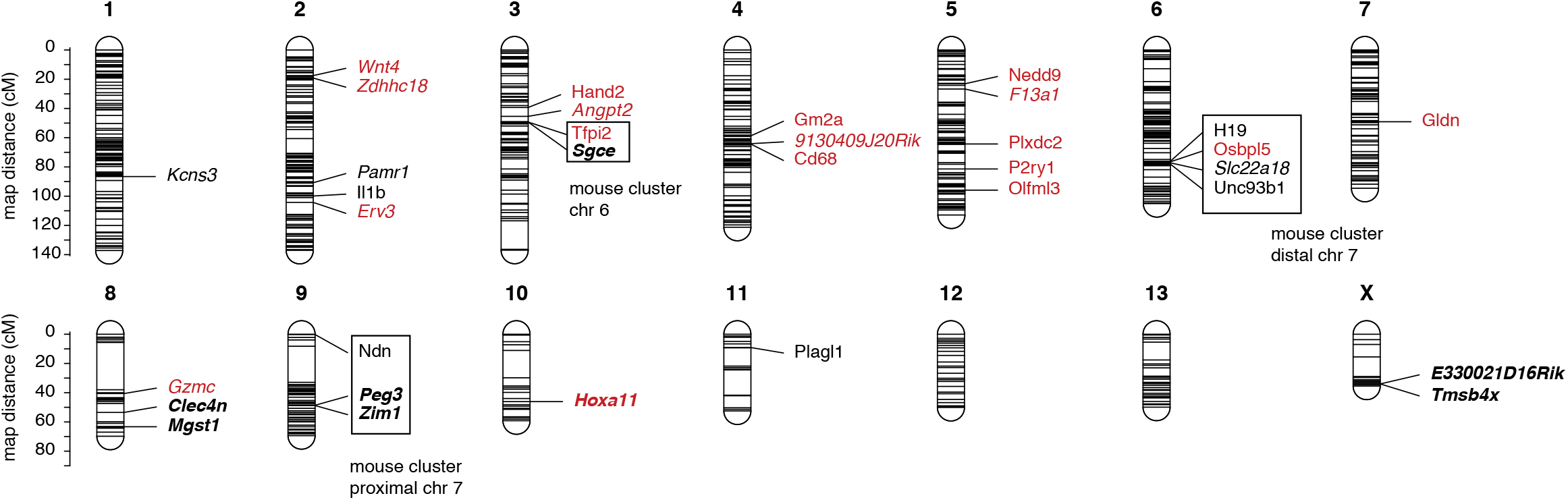
Candidate imprinted genes placed on the genetic map. Candidate imprinted genes in the BC downregulated module are indicated in red, and clusters with potential homology to imprinted clusters in *Mus* are indicated with boxes. Visualized with R/LinkageMapView (Ouellette et al. 2018).

**Figure S15.**
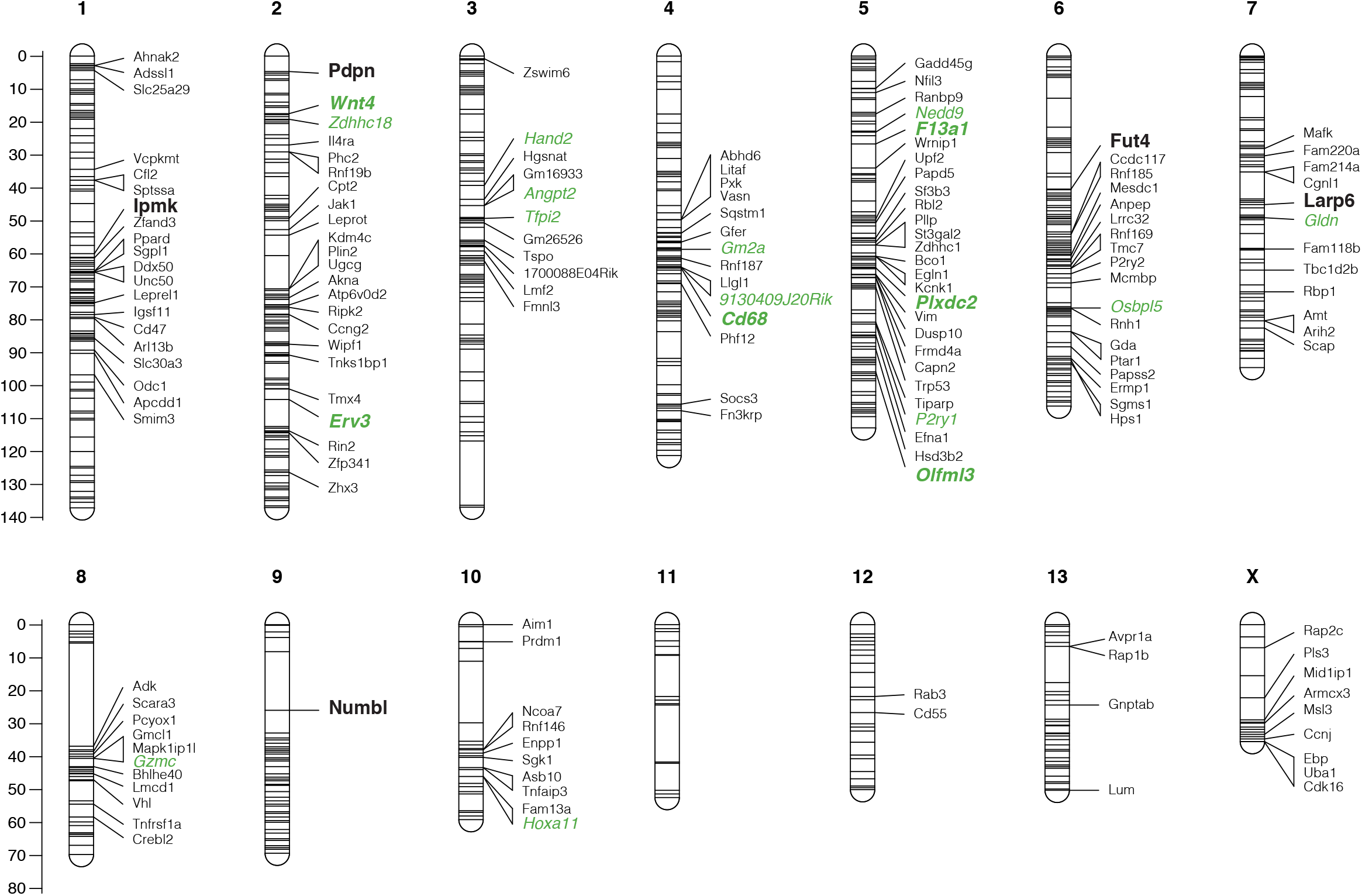
BC downregulated module placed on the genetic map. BC downregulated network hub genes are in bold, and candidate imprinted are indicated in green. Visualized with R/LinkageMapView (Ouellette et al. 2018)

**Figure S16.**
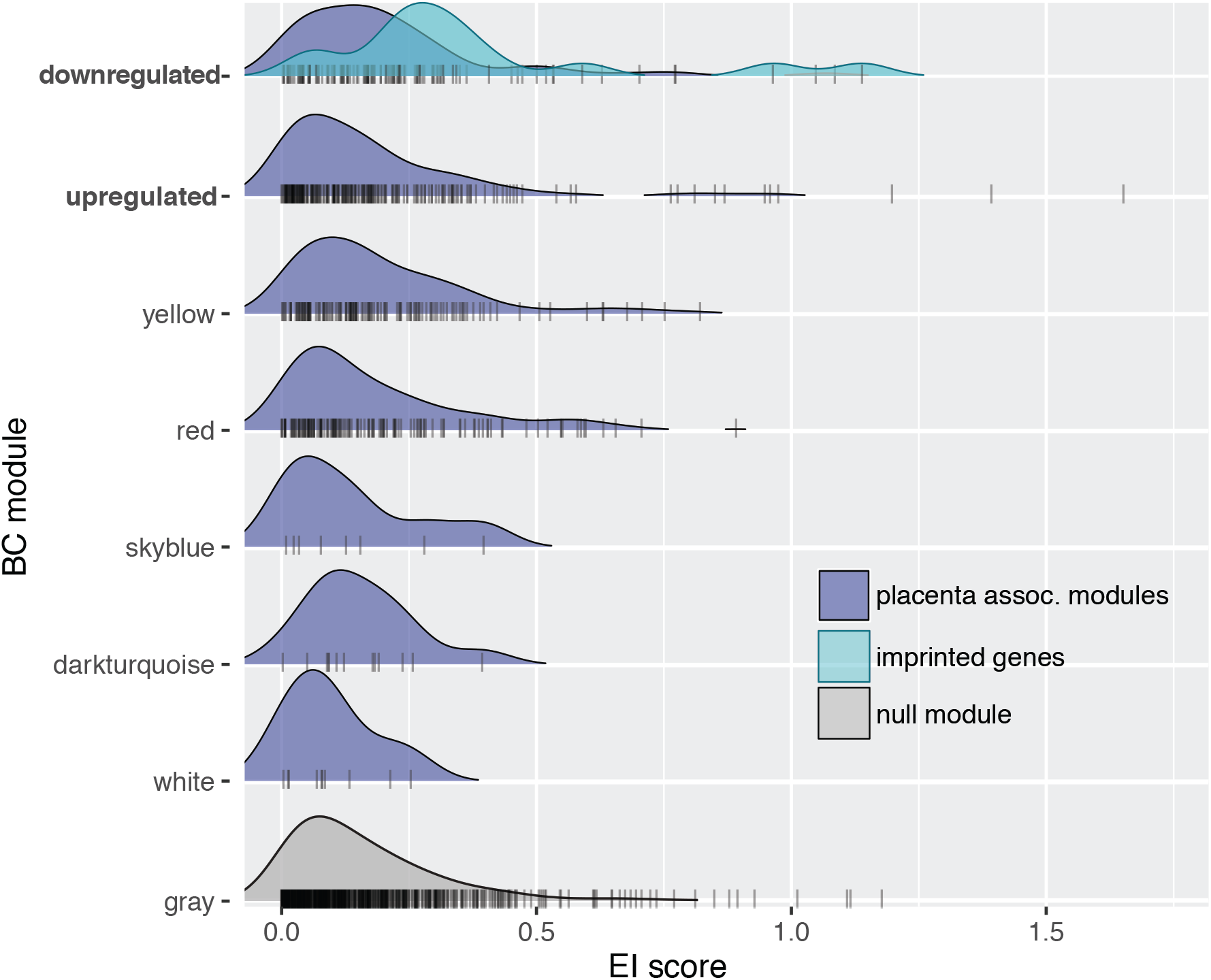
EI score distributions by module and imprinting status. The gray module includes all genes that were not placed in any module in the network analysis, and serves as the null expectation of the distribution of the score. All BC modules associated with placenta size are shown. Candidate imprinted genes in the downregulated module showed a shift towards increased EI values, while other placenta associated modules did not.

**Figure S17.**
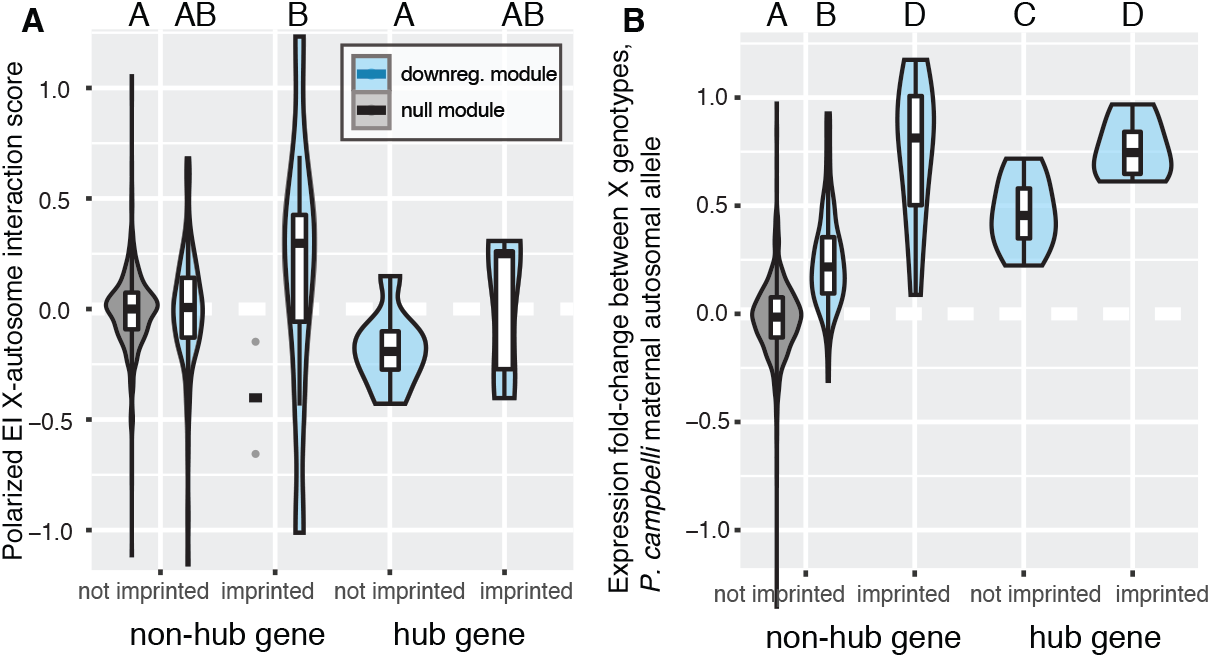
Polarized EI score distributions by hub and imprinting status. (A) Polarized expression interaction scores indicated that imprinted genes were more likely to have a larger fold change when maternal alleles were mismatched with the maternal X chromosome (a positive value). (B) Positive values for gene expression fold change between X genotypes for individuals with a (mismatched) homozygous *P. campbelli* autosomal genotype indicated that expression was higher for individuals with a *P. campbelli* X chromosome than for those with a *P. sungorus* X chromosome. All letter groups indicate significance based on a Tukey’s HSD test, P < 0.05. and imprinting status. The grey module includes all genes that were not placed in any module in the network analysis, and serves as the null expectation of the distribution of the score. All BC modules associated with placenta size are included here, with shift of increased EI values for the candidate imprinted genes in the downregulated module, but not the other placenta associated modules.

**Figure S18.**
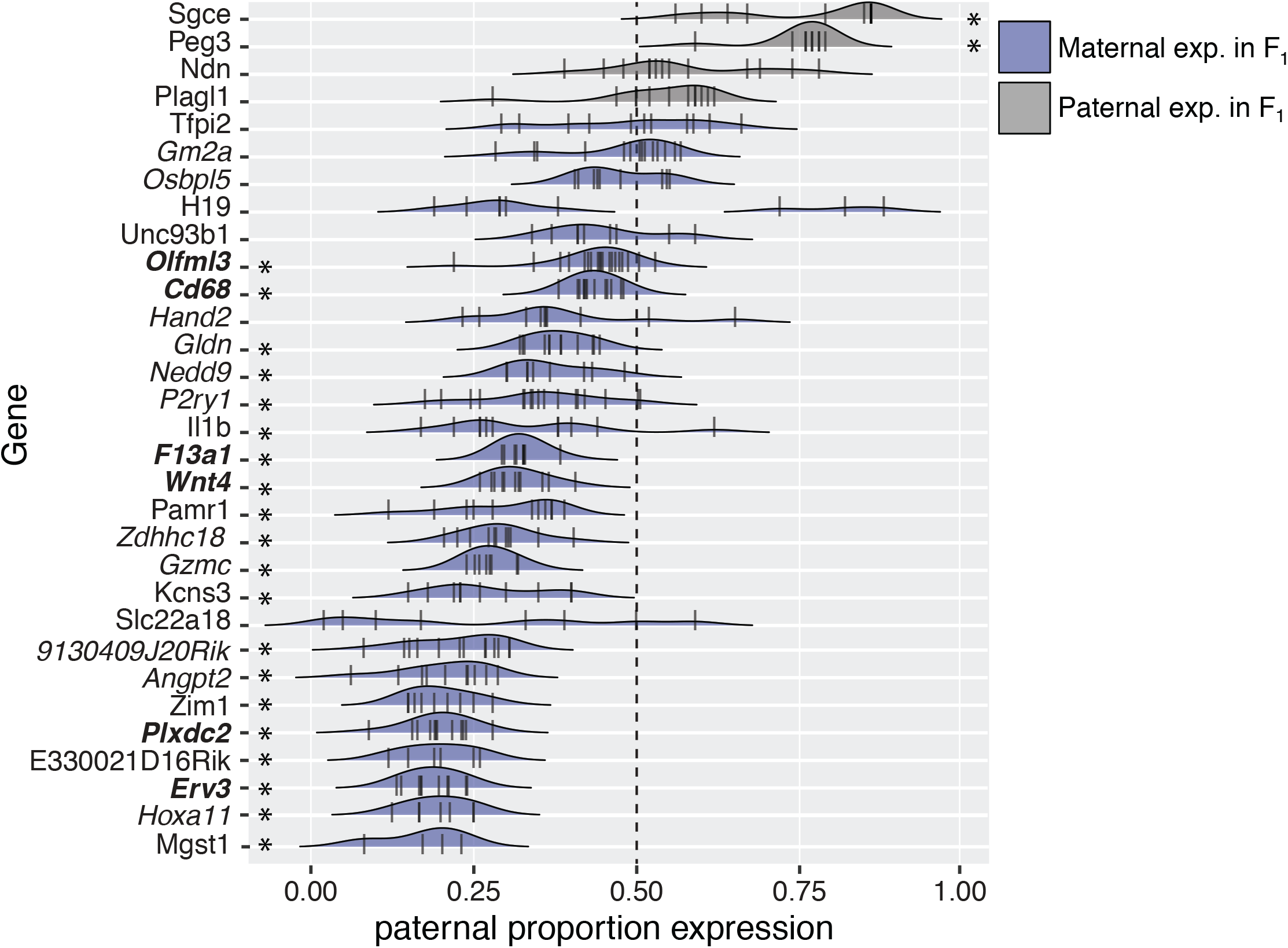
Allelic bias in candidate imprinted genes in BC. Proportion of paternal expression for all heterozygous BC hybrids for each of the 31 candidate imprinted genes that were placed on the map. To calculate allele-specific expression, the reference (*P. sungorus*) allele was designated as the maternal allele and the proportion of maternal reads was averaged over variable sites for heterozygous BC individuals, excluding all homozygous individuals at each gene. Generally, we recover qualitatively consistent signals for allelic bias at genes that displayed maternal (blue) or paternal bias (gray) in F_1_ hybrids. Bolded genes are hub genes in the BC downregulated module. Stars indicate that mean paternal expression was significantly different from 0.5 (one-sample T-test, Bonferroni corrected P < 0.05).

**Figure S19.**
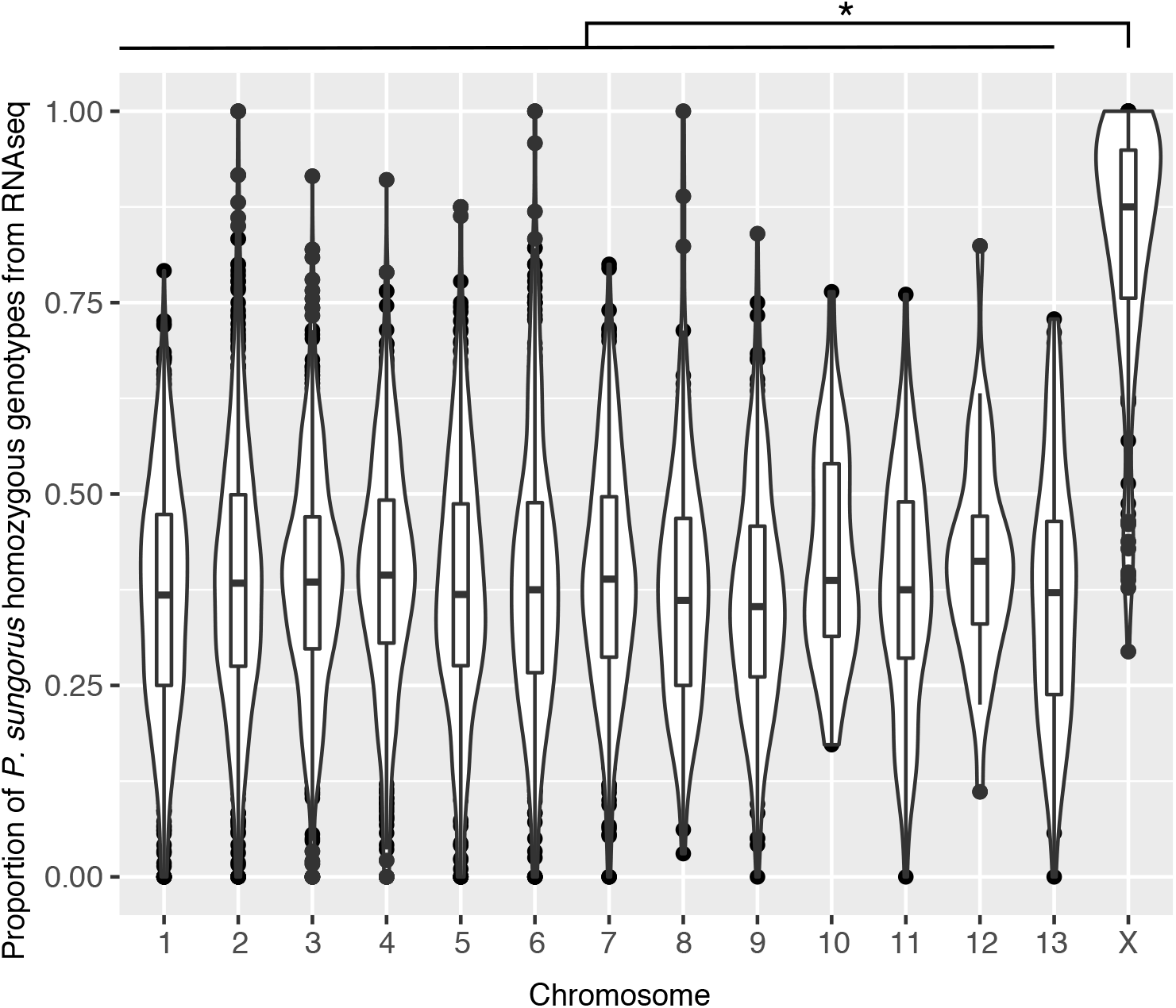
Allelic bias on the X chromosome in heterozygous BC females. The average proportion of genotypes for each mapped gene called as homozygous for the maternal *P. sungorus* allele is shown for each chromosome. Estimates for each gene were based on RNAseq reads from females inferred to be heterozygous based on their local genetic map genotype. No backcross individuals should be homozygous for the *P. sungorus* allele based on standard diploid expectations and unbiased gene expression, while paternally imprinted XCI should generate only homozygous *P. sungorus* genotypes in heterozygous females (proportion = 1.0). The autosomes showed a skew towards expression of *P. sungorus* alleles relative to these predictions, likely reflecting errors in the genetic map genotyping, calling heterozygous genotypes from using RNAseq data, and uncertainty in species origin of individual variants. Overall, the X chromosome showed strong allelic bias towards the *P. sungorus* maternal allele, consistent with intact paternal XCI. Star indicates a significant difference (adjusted r^2^ = 0.17, *F*_1,3560_ = 930.3, P << 0.0001, ANOVA).

**Table S1.** A full description of all RAD markers including their ID, the linkage group they are found on, genetic position in centiMorgans, the position and polarization of the diagnostic SNV between *P. campbelli* and *P. sungorus*, and the sequence of the marker which always begin with TGCAGG (the restriction-enzyme cut-site of SbfI, i.e.: CC_TGCA^GG). SNVs in the sequence are denoted with standard IUPAC ambiguity codes.

**Table S2.** WGCNA modules generated from F_1_ and pure species placental gene expression data. Color names are arbitrarily and randomly generated by the program, and have no additional meaning. The upregulated and downregulated modules are discussed in the manuscript are indicated as such. Counts of genes in each module, correspondence with previous pairwise analysis (Brekke *et al*. 2016), association with inheritance pattern and phenotypes, and enrichment for candidate imprinted genes indicated.

**Table S3.** WGCNA modules generated from BC placental gene expression data. As before, color names are arbitrarily and randomly generated by the program, and do not correspond whatsoever with the arbitrarily assigned names given to the F_1_ data. The upregulated and downregulated modules are discussed in the manuscript are indicated as such. Counts of genes in each module, correspondence with F_1_ network analysis, association with phenotypes, and enrichment for candidate imprinted genes indicated.

**Table S4. A full description of the genetic locations of each gene from that was captured and associated with the map.** Columns are: Linkage group (LG), position in centiMorgans (cM), gene name from the *P. sungorus* transcriptome (Trinity_Component), the exon that the SNP appears in (exon), the position of the SNP in the exon (snp_pos_in_exon), the gene name (Associated_Gene_Name), and the mouse ensemble gene ID of that gene (Ensembl_Gene_ID).

**Table S5.** SRA sequence accession numbers for each individual by sequence type.

## References

Al Adhami, H., B. Evano, A. Le Digarcher, C. Gueydan, E. Dubois et al., 2015 A systems-level approach to parental genomic Imprinting: the imprinted gene network includes extracellular matrix genes and regulates cell cycle exit and differentiation. Genome Res. 25: 353–367.

Allen, W. R., J. A. Skidmore, F. Stewart and D. F. Antczak, 1993 Effects of fetal genotype and uterine environment on placental development in equids. J. Reprod. Fertil. 98: 55–60.

Arévalo, L., S. Gardner and P. Campbell, 2020 Haldane’s rule in the placenta: sex-biased misregulation of the Kcnq1 imprinting cluster in hybrid mice. bioRxiv doi: 10.1101/2020.05.07.082248.

Astarita, J. L., S. E. Acton and S. J. Turley, 2012 Podoplanin: emerging functions in development the immune system and cancer. Front. Immunol. 3: 283.

Babak, T., B. DeVeale, E. K. Tsang, Y. Q. Zhou, X. Li et al., 2015 Genetic conflict reflected in tissue-specific maps of genomic imprinting in human and mouse. Nat. Genet. 47: 544–549.

Bao, R., K. G. Onishi, E. Tolla, F. J. P. Ebling, J. E. Lewis et al., 2019 Genome sequencing and transcriptome analyses of the Siberian hamster hypothalamus identify mechanisms for seasonal energy balance. Proc. Natl. Acad. Sci. USA 116: 13116–13121.

Bateson, W., 1909 Heredity and variation in modern lights., pp. 85-101 in Darwin and Modern Science, edited by A. C. Seward. Cambridge University Press, Cambridge.

Bhattacharyya, T., S. Gregorova, O. Mihola, M. Anger, J. Sebestova et al., 2013 Mechanistic basis of infertility of mouse intersubspecific hybrids. Proc. Natl. Acad. Sci. USA 110: E468–E477.

Bikchurina, T. I., K. V. Tishakova, E. A. Kizilova, S. A. Romanenko, N. A. Serdyukova et al., 2018 Chromosome synapsis and recombination in male-sterile and female-fertile interspecies hybrids of the dwarf hamsters (Phodopus, Cricetidae). Genes 9: 27.

Bolger, A. M., M. Lohse and B. Usadel, 2014 Trimmomatic: a flexible trimmer for Illumina sequence data. Bioinformatics 30: 2114–2120.

Bouwens, E. A. M., F. Stavenuiter and L. O. Mosnier, 2013 Mechanisms of anticoagulant and cytoprotective actions of the protein C pathway. J. Thromb. Haemost. 11: 242–253.

Brekke, T., L. Henry and J. Good, 2016 Genomic imprinting, disrupted placental expression, and speciation. Evolution 70: 2690–2703.

Brekke, T. D., and J. M. Good, 2014 Parent-of-origin growth effects and the evolution of hybrid inviability in dwarf hamsters. Evolution 68: 3134–3148.

Brekke, T. D., K. A. Steele and J. F. Mulley, 2018 Inbred or outbred? Genetic diversity in laboratory rodent colonies. G3 8: 679–686.

Brekke, T. D., S. Supriya, M. G. Denver, A. Thom, K. A. Steele et al., 2019 A high-density genetic map and molecular sex-typing assay for gerbils. Mamm. Genome 30: 63–70.

Broman, K. W., and S. Sen, 2009 A guide to QTL mapping with R/qtl. Springer, New York.

Brown, J. D., V. Piccuillo and R. J. O’Neill, 2012 Retroelement demethylation associated with abnormal placentation in Mus musculus x Mus caroli hybrids. Biol. Reprod. 86: 88–88.

Burgin, C. J., J. P. Colella, P. L. Kahn and N. S. Upham, 2018 How many species of mammals are there? J. Mammal. 99: 1–14.

Butlin, R., A. Debelle, C. Kerth, R. R. Snook, L. W. Beukeboom et al., 2012 What do we need to know about speciation? Trends Ecol. Evol. 27: 27–39.

Campbell, P., J. M. Good and M. D. Nachman, 2013 Meiotic sex chromosome inactivation is disrupted in sterile hybrid male house mice. Genetics 193: 819–828.

Capellini, I., C. Venditti and R. A. Barton, 2011 Placentation and maternal investment in mammals. Am. Nat. 177: 86–98.

Carroll, S. B., 2008 Evo-devo and an expanding evolutionary synthesis: A genetic theory of morphological evolution. Cell 134: 25–36.

Catchen, J., P. A. Hohenlohe, S. Bassham, A. Amores and W. A. Cresko, 2013 Stacks: an analysis tool set for population genomics. Mol. Ecol. 22: 3124–3140.

Chen, Z. Y., D. E. Hagen, C. G. Elsik, T. Ji, C. J. Morris et al., 2015 Characterization of global loss of imprinting in fetal overgrowth syndrome induced by assisted reproduction. Proc. Natl. Acad. Sci. USA 112: 4618–4623.

Chen, Z. Y., D. E. Hagen, J. B. Wang, C. G. Elsik, T. M. Ji et al., 2016 Global assessment of imprinted gene expression in the bovine conceptus by next generation sequencing. Epigenetics 11: 501–516.

Cheng, G., M. Zhong, R. Kawaguchi, M. Kassai, M. Al-Ubaidi et al., 2014 Identification of PLXDC1 and PLXDC2 as the transmembrane receptors for the multifunctional factor PEDF. eLife 3: e05401.

Chistiakov, D. A., M. C. Killingsworth, V. A. Myasoedova, A. N. Orekhov and Y. V. Bobryshev, 2017 CD68/macrosialin: not just a histochemical marker. Lab. Invest. 97: 4–13.

Chuong, E. B., 2018 The placenta goes viral: Retroviruses control gene expression in pregnancy. PLoS Biol. 16: e3000028.

Cochery-Nouvellon, E., C. Chauleur, C. Demattei, E. Mercier, P. Fab-bro-Peray et al., 2009 The A6936G polymorphism of the endothelial protein C receptor gene is associated with the risk of unexplained foetal loss in Mediterranean European couples. Thromb. Haemost. 102: 656–667.

Constancia, M., M. Hemberger, J. Hughes, W. Dean, A. Ferguson-Smith et al., 2002 Placental-specific IGF-II is a major modulator of placental and fetal growth. Nature 417: 945–948.

Coughlan, J. M., M. W. Brown and J. H. Willis, 2020 Patterns of hybrid seed inviability in the Mimulus guttatus sp. complex reveal a potential role of parental conflict in reproductive isolation. Curr. Biol. 30: 83–93.

Coughlan, J. M., and D. R. Matute, 2020 The importance of intrinsic post-zygotic barriers throughout the speciation process. Philos. Trans. R. Soc. Lond., B, Biol. Sci. 375.

Coyne, J. A., and H. A. Orr, 1989a Patterns of speciation in Drosophila. Evolution 43: 362–381.

Coyne, J. A., and H. A. Orr, 1989b Two rules of speciation., pp. 180–207 in Speciation and Its Consequences, edited by D. Otte and J. Endler. Sinauer Associates, Sunderland, MA.

Coyne, J. A., and H. A. Orr, 2004 Speciation. Sinauer Associates, Inc., Sunderland, MA.

Crespi, B., and P. Nosil, 2013 Conflictual speciation: species formation via genomic conflict. Trends Ecol. Evol. 28: 48–57.

Davis, B. W., C. M. Seabury, W. A. Brashear, G. Li, M. Roelke-Parker et al., 2015 Mechanisms underlying mammalian hybrid sterility in two feline interspecies models. Mol. Biol. Evol. 32: 2534–2546.

Dobzhansky, T., 1937 Genetics and the Origin of Species. Columbia University Press, New York.

Dupont, C., and J. Gribnau, 2013 Different flavors of X-chromosome inactivation in mammals. Curr. Opin. Cell Biol. 25: 314–321.

Ebina, Y., M. Ieko, S. Naito, G. Kobashi, M. Deguchi et al., 2015 Low levels of plasma protein S, protein C and coagulation factor XII during early pregnancy and adverse pregnancy outcome. Thromb. Haemost. 114: 65–69.

Feenstra, B., I. M. Skovgaard and K. W. Broman, 2006 Mapping quantitative trait loci by an extension of the Haley-Knott regression method using estimating equations. Genetics 173: 2269–2282.

Ferreira, D. M. S., A. J. Cheng, L. Z. Agudelo, I. Cervenka, T. Chaillou et al., 2019 LIM and cysteine-rich domains 1 (LMCD1) regulates skeletal muscle hypertrophy, calcium handling, and force. Skelet. Muscle 9: 26.

Finn, E. H., C. L. Smith, J. Rodriguez, A. Sidow and J. C. Baker, 2014 Maternal bias and escape from X chromosome imprinting in the midgestation mouse placenta. Dev. Biol. 390: 80–92.

Gamperl, R., G. Vistorin and W. Rosenkranz, 1977 New observations on karyotype of Djungarian hamster, Phodopus sungorus. Experientia 33: 1020–1021.

Garner, A. G., A. M. Kenney, L. Fishman and A. L. Sweigart, 2016 Genetic loci with parent-of-origin effects cause hybrid seed lethality in crosses between Mimulus species. New Phytol. 211: 319–331.

Ghazalpour, A., S. Doss, B. Zhang, S. Wang, C. Plaisier et al., 2006 Integrating genetic and network analysis to characterize genes related to mouse weight. PLoS Genet. 2: 1182–1192.

Good, J. M., M. D. Dean and M. W. Nachman, 2008 A complex genetic basis to X-linked hybrid male sterility between two species of house mice. Genetics 179: 2213–2228.

Gris, J. C., S. Bouvier, E. Cochery-Nouvellon, E. Mercier, E. Mousty et al., 2019 The role of haemostasis in placenta-mediated complications. Thromb. Res. 181: S10–S14.

Guerrero, R. F., A. L. Posto, L. C. Moyle and M. W. Hahn, 2016 Ge-nome-wide patterns of regulatory divergence revealed by introgression lines. Evolution 70: 696–706.

Haaf, T., H. Weis and M. Schmid, 1987 A comparative cytogenetic study on the mitotic and meiotic chromosomes in hamster species of the genus Phodopus (Rodentia, Cricetinae). Z. Säugetierkd. 52: 281–290.

Haig, D., 1996 Placental hormones, genomic imprinting, and maternal-fetal communication. J. Evol. Biol. 9: 357–380.

Haig, D., 2000 The kinship theory of genomic imprinting. Annu. Rev. Ecol. Syst. 31: 9–32.

Haley, C. S., and S. A. Knott, 1992 A simple regression method for mapping quantitative trait loci in line crosses using flanking markers. Heredity 69: 315–324.

Heard, E., and C. M. Disteche, 2006 Dosage compensation in mammals: fine-tuning the expression of the X chromosome. Gene Dev. 20: 1848–1867.

Hemberger, M., 2002 The role of the X chromosome in mammalian extra embryonic development. Cytogenetic and Genome Research 99: 210–217.

Hemberger, M. C., R. S. Pearsall, U. Zechner, A. Orth, S. Otto et al., 1999 Genetic dissection of X-linked interspecific hybrid placental dysplasia in congenic mouse strains. Genetics 153: 383–390.

Hirasawa, R., and R. Feil, 2010 Genomic imprinting and human disease, pp. 187-200 in Essays in Biochemistry: Epigenetics, Disease and Behaviour, edited by H. J. Lipps, J. Postberg and D. A. Jackson.

Hudson, Q. J., T. M. Kulinski, S. P. Huetter and D. P. Barlow, 2010 Genomic imprinting mechanisms in embryonic and extraembryonic mouse tissues. Heredity 105: 45–56.

Isermann, B., R. Sood, R. Pawlinski, M. Zogg, S. Kalloway et al., 2003 The thrombomodulin-protein C system is essential for the maintenance of pregnancy. Nat. Med. 9: 331–337.

Ishishita, S., K. Tsuboi, N. Ohishi, K. Tsuchiya and Y. Matsuda, 2015 Abnormal pairing of X and Y sex chromosomes during meiosis I in interspecific hybrids of Phodopus campbelli and P. sungorus. Sci. Rep. 5: 9435–9439.

Jang, S., D. Oh, Y. Lee, E. Hosy, H. Shin et al., 2016 Synaptic adhesion molecule IgSF11 regulates synaptic transmission and plasticity. Nat. Neurosci. 19: 84–93.

Jin, M., K. Udagawa, E. Miyagi, T. Nakazawa, F. Hirahara et al., 2001 Expression of serine proteinase inhibitor PP5/TFPI-2/MSPI decreases the invasive potential of human choriocarcinoma cells in vitro and in vivo. Gynecol. Oncol. 83: 325–333.

Kaneko-Ishino, T., and F. Ishino, 2019 Evolution of viviparity in mammals: what genomic imprinting tells us about mammalian placental evolution. Reprod. Fertil. Dev. 31: 1219–1227.

Karabag, T., A. Kaya, A. Temizhan, F. Koc, S. Yavuz et al., 2007 The influence of homocysteine levels on endothelial function and their relation with microvascular complications in T2DM patients without macrovascular disease. Acta Diabetol. 44: 69–75.

Kent, W. J., 2002 BLAT - The BLAST-like alignment tool. Genome Res. 12: 656–664.

Khil, P. P., N. A. Smirnova, P. J. Romanienko and R. D. Camerini-Otero, 2004 The mouse X chromosome is enriched for sex-biased genes not subject to selection by meiotic sex chromosome inactivation. Nat. Genet. 36: 642–646.

King, M. C., and A. C. Wilson, 1975 Evolution at two levels in humans and chimpanzees. Science 188: 107–116.

Knoefler, M., and J. Pollheimer, 2013 Human placental trophoblast invasion and differentiation: a particular focus on Wnt signaling. Front. Genet. 4: 190.

Konduri, S. D., A. Tasiou, N. Chandrasekar and J. S. Rao, 2001 Over-expression of tissue factor pathway inhibitor-2 (TFPI-2), decreases the invasiveness of prostate cancer cells in vitro. Int. J. Oncol. 18: 127–131.

Kurz, H., U. Zechner, A. Orth and R. Fundele, 1999 Lack of correlation between placenta and offspring size in mouse interspecific crosses. Anat. Embryol. 200: 335–343.

Langfelder, P., and S. Horvath, 2008 WGCNA: an R package for weighted correlation network analysis. BMC Bioinform. 9: 559.

Langmead, B., and S. L. Salzberg, 2012 Fast gapped-read alignment with Bowtie 2. Nat. Methods 9: 357–359.

Larson, E., D. Vanderpool, S. Keeble, M. Dean and J. Good, 2017 The composite regulatory basis of the large X-effect in mouse speciation. Mol. Biol. Evol. 34: 282–295.

Larson, E. L., E. E. K. Kopania and J. M. Good, 2018 Spermatogenesis and the evolution of mammalian sex chromosomes. Trends Genet. 34: 722–732.

Lee, J. T., and M. S. Bartolomei, 2013 X-inactivation, imprinting, and long noncoding RNAs in health and disease. Cell 152: 1308–1323.

Levy, O., 2007 Innate immunity of the newborn: basic mechanisms and clinical correlates. Nat. Rev. Immunol. 7: 379–390.

Li, H., and R. Durbin, 2009 Fast and accurate short read alignment with Burrows-Wheeler transform. Bioinformatics 25: 1754–1760.

Li, J. Y., N. Paragas, R. M. Ned, A. Qiu, M. Viltard et al., 2009 Scara5 Is a ferritin receptor mediating non-transferrin iron delivery. Dev. Cell 16: 35–46.

Li, M., N. M. J. Schwerbrock, P. M. Lenhart, K. L. Fritz-Six, M. Kadmiel et al., 2013 Fetal-derived adrenomedullin mediates the innate immune milieu of the placenta. J. Clin. Investig. 123: 2408–2420.

Liao, Y., G. K. Smyth and W. Shi, 2014 featureCounts: an efficient general purpose program for assigning sequence reads to genomic features. Bioinformatics 30: 923–930.

Lifschytz, E., and D. L. Lindsley, 1972 The role of X-chromosome inactivation during spermatogenesis. Proc. Natl. Acad. Sci. USA 69: 182–186.

Loschiavo, M., Q. K. Nguyen, A. R. Duselis and P. B. Vrana, 2007 Mapping and identification of candidate loci responsible for Peromyscus hybrid overgrowth. Mamm. Genome 18: 75–85.

Lyon, M. F., 1961 Gene action in X-chromosome of mouse (Mus musculus L). Nature 190: 372–373.

Mack, K. L., M. Phifer-Rixey, B. Harr and M. W. Nachman, 2019 Gene expression networks across multiple tissues are associated with rates of molecular evolution in wild house mice. Genes 10: 225.

Malabanan, M. M., and R. D. Blind, 2016 Inositol polyphosphate multikinase (IPMK) in transcriptional regulation and nuclear inositide metabolism. Biochem. Soc. Trans. 44: 279–285.

Mangeney, M., M. Renard, G. Schlecht-Louf, I. Bouallaga, O. Heidmann et al., 2007 Placental syncytins: Genetic disjunction between the fusogenic and immunosuppressive activity of retroviral envelope proteins. Proc. Natl. Acad. Sci. USA 104: 20534–20539.

Manshouri, R., E. Coyaud, S. T. Kundu, D. H. Peng, S. A. Stratton et al., 2019 ZEB1/NuRD complex suppresses TBC1D2b to stimulate E-cadherin internalization and promote metastasis in lung cancer. Nat. Comm. 10: 5125.

Martin, M., 2011 Cutadapt removes adapter sequences from high-throughput sequencing reads. EMBnet J. 17: 1–3.

Masly, J. P., and D. C. Presgraves, 2007 High-resolution genome-wide dissection of the two rules of speciation in Drosophila. PLoS Biol. 5: 1890–1898.

McGraw, S., C. C. Oakes, J. Martel, M. C. Cirio, P. de Zeeuw et al., 2013 Loss of DNMT1o disrupts imprinted X chromosome inactivation and accentuates placental defects in females. PLoS Genet. 9.

Meyer, M., and M. Kircher, 2010 Illumina sequencing library preparation for highly multiplexed target capture and sequencing. Cold Spring Harb. Protoc. 2010: ppdb.prot5448.

Miljkovic-Licina, M., P. Hammel, S. Garrido-Urbani, B. P. L. Lee, M. Megue-nani et al., 2012 Targeting Olfactomedin-like 3 inhibits tumor growth by impairing angiogenesis and pericyte coverage. Mol. Cancer Ther. 11: 2588–2599.

Miller, S. F. C., K. Summerhurst, A. E. Runker, G. Kerjan, R. H. Friedel et al., 2007 Expression of Plxdc2/TEM7R in the developing nervous system of the mouse. Gene Expr. Patterns 7: 635–644.

Moreira de Mello, J. C., E. S. Souza de Araujo, R. Stabellini, A. M. Fraga, J. E. Santana de Souza et al., 2010 Random X inactivation and extensive mosaicism in human placenta revealed by analysis of allele-specific gene expression along the X chromosome. PLoS One 5: e10947.

Morgan, K., B. Harr, M. A. White, B. A. Payseur and L. M. Turner, 2020 Disrupted gene networks in subfertile hybrid house mice. Mol. Biol. Evol. 37: 1547–1562.

Morison, I. M., J. P. Ramsay and H. G. Spencer, 2005 A census of mammalian imprinting. Trends Genet. 21: 457–465.

Muller, H. J., 1942 Isolating mechanisms, evolution, and temperature. Biol. Symp. 6: 71–125.

Muszbek, L., Z. Bereczky, Z. Bagoly, I. Komaromi and E. Katona, 2011 Factor XIII: a coagulation factor with multiple plasmatic and cellular functions. Physiol. Rev. 91: 931–972.

O’Brien, E. K., and J. B. Wolf, 2017 The coadaptation theory for genomic imprinting. Evol. Lett. 1: 49–59.

Okamoto, I., C. Patrat, D. Thepot, N. Peynot, P. Fauque et al., 2011 Eutherian mammals use diverse strategies to initiate X-chromosome inactivation during development. Nature 472: 370–374.

Orr, H. A., and M. Turelli, 2001 The evolution of postzygotic isolation: accumulating Dobzhansky-Muller incompatibilities. Evolution 55: 1085–1094.

Ouellette, L. A., R. W. Reid, S. G. Blanchard and C. R. Brouwer, 2018 LinkageMapView-rendering high-resolution linkage and QTL maps. Bioinformatics 34: 306–307.

Patten, M. M., M. Cowley, R. J. Oakey and R. Feil, 2016 Regulatory links between imprinted genes: evolutionary predictions and consequences. P. R. Soc. B 283: 20152760.

Pavlicev, M., K. Hiratsuka, K. A. Swaggart, C. Dunn and L. Muglia, 2015 Detecting endogenous retrovirus-driven tissue-specific gene transcription. Genome Biol. Evol. 7: 1082–1097.

Peterson, B. K., J. N. Weber, E. H. Kay, H. S. Fisher and H. E. Hoekstra, 2012 Double digest RADseq: an inexpensive method for de novo SNP discovery and genotyping in model and non-model species. PLoS One 7: e37135.

Plasschaert, R. N., and M. S. Bartolomei, 2014 Genomic imprinting in development, growth, behavior and stem cells. Development 141: 1805–1813.

Reik, W., M. Constancia, A. Fowden, N. Anderson, W. Dean et al., 2003 Regulation of supply and demand for maternal nutrients in mammals by imprinted genes. J. Physiol. 547: 35–44.

Ribarska, T., M. Ingenwerth, W. Goering, R. Engers and W. A. Schulz, 2010 Epigenetic inactivation of the placentally imprinted tumor suppressor gene TFPI2 in prostate carcinoma. Cancer Genomics Proteomics 7: 51–60.

Robinson, M. D., and A. Oshlack, 2010 A scaling normalization method for differential expression analysis of RNA-seq data. Genome Biol. 11: R25.

Romanenko, S. A., V. T. Volobouev, P. L. Perelman, V. S. Lebedev, N. A. Serdukova et al., 2007 Karyotype evolution and phylogenetic relationships of hamsters (Cricetidae, Muroidea, Rodentia) inferred from chromosomal painting and banding comparison. Chromosome Res. 15: 283–297.

Safronova, L. D., E. V. Cherepanova and N. Y. Vasil’eva, 1999 Specific features of the first meiotic division in hamster hybrids obtained by backcrossing Phodopus sungorus and Phodopus campbelli. Russ. J. Genet. 35: 184–188.

Sanli, I., and R. Feil, 2015 Chromatin mechanisms in the developmental control of imprinted gene expression. Int. J. Biochem. 67: 139–147.

Schmidt, A., N. Endo, S. J. Rutledge, R. Vogel, D. Shinar et al., 1992 Identification of a new member of the steroid-hormone receptor superfamily that is activated by a peroxisome proliferator and fatty-acids. Mol. Endocrinol. 6: 1634–1641.

Scribner, S. J., and K. E. Wynne-Edwards, 1994 Disruption of body-temperature and behavior rhythms during reproduction in dwarf hamster (Phodopus). Physiology & Behavior 55: 361–369.

Sears, K. E., J. A. Maier, M. Rivas-Astroza, R. Poe, S. Zhong et al., 2015 The relationship between gene network structure and expression variation among individuals and species. PLoS Genet. 11: e1005398.

Shi, W., L. Lefebvre, Y. Yu, S. Otto, A. Krella et al., 2004 Loss-of-imprinting of Peg1 in mouse interspecies hybrids is correlated with altered growth. genesis 39: 65–72.

Sonderegger, S., J. Pollheimer and M. Knoefler, 2010 Wnt signalling in implantation, decidualisation and placental Differentiation - Review. Placenta 31: 839–847.

Sood, R., S. Kalloway, A. E. Mast, C. J. Hillard and H. Weiler, 2006 Feto-maternal cross talk in the placental vascular bed: control of coagulation by trophoblast cells. Blood 107: 3173–3180.

Stefanovic, B., Z. Manojlovic, C. Vied, C. D. Badger and L. Stefanovic, 2019 Discovery and evaluation of inhibitor of LARP6 as specific antifibrotic compound. Sci. Rep. 9: 326.

Storchová, R., S. Gregorová, D. Buckiová, V. Kyselová, P. Divina et al., 2004 Genetic analysis of X-linked hybrid sterility in the house mouse. Mamm. Genome 15: 515–524.

Storey, J. D., A. J. Bass, A. Dabney and D. Robinson, 2019 qvalue: Q-value estimation for false discovery rate control., pp. R package version 2.18.0.

Takagi, N., and M. Sasaki, 1975 Preferential inactivation of paternally derived X-chromosome in extraembryonic membranes of mouse. Nature 256: 640–642.

Theiler, K., 1972 The house mouse: Development and normal stages from fertilization to 4 weeks of age. Springer-Verlag, New York, NY.

Turelli, M., and L. C. Moyle, 2007 Asymmetric postmating isolation: Dar-win’s corallary to Haldane’s rule. Genetics 176: 1059–1088.

Turelli, M., and H. A. Orr, 2000 Dominance, epistasis and the genetics of postzygotic isolation. Genetics 154: 1663–1679.

Van der Auwera, G. A., M. O. Carneiro, C. Hartl, R. Poplin, G. del Angel et al., 2013 From FastQ data to high confidence variant calls: the Genome Analysis Toolkit best practices pipeline. Curr. Protoc. Bioinformatics 43: 11–33.

Varrault, A., C. Gueydan, A. Delalbre, A. Bellmann, S. Houssami et al., 2006 Zac1 regulates an imprinted gene network critically involved in the control of embryonic growth. Dev. Cell 11: 711–722.

Vrana, P. B., 2007 Genomic imprinting as a mechanism of reproductive isolation in mammals. J. Mammal. 88: 5–23.

Vrana, P. B., J. A. Fossella, P. Matteson, T. del Rio, M. J. O’Neill et al., 2000 Genetic and epigenetic incompatibilities underlie hybrid dysgenesis in Peromyscus. Nat. Genet. 25: 120–124.

Wang, H., M. Morales-Levy, J. Rose, L. C. Mackey, P. Bodary et al., 2013 α(1,3)-Fucosyltransferases FUT4 and FUT7 control murine susceptibility to thrombosis. Am. J. Pathol. 182: 2082–2093.

Wang, X., D. C. Miller, A. G. Clark and D. F. Antczak, 2012 Random X inactivation in the mule and horse placenta. Genome Res. 22: 1855–1863.

Wang, X., P. D. Soloway and A. G. Clark, 2011 A survey for novel imprinted genes in the mouse placenta by mRNA-seq. Genetics 189: 109–122.

Wang, Z., M. Gerstein and M. Snyder, 2009 RNA-Seq: a revolutionary tool for transcriptomics. Nat. Rev. Genet. 10: 57–63.

Wilder, J. A., E. K. Hewett and M. E. Gansner, 2009 Molecular evolution of GYPC: Evidence for recent structural innovation and positive selection in humans. Mol. Biol. Evol. 26: 2679–2687.

Wilson, A., D. L. Ardiet, C. Saner, N. Vilain, F. Beermann et al., 2007 Normal hemopoiesis and lymphopoiesis in the combined absence of numb and numblike. J. Immunol. 178: 6746–6751.

Wolf, J. B., 2013 Evolution of genomic imprinting as a coordinator of coadapted gene expression. Proc. Natl. Acad. Sci. USA 110: 5085–5090.

Wolf, J. B., and Y. Brandvain, 2014 Gene interactions in the evolution of genomic imprinting. Heredity 113: 129–137.

Wolf, J. B., and R. Hager, 2006 A maternal–offspring coadaptation theory for the evolution of genomic imprinting. PLoS Biol. 4: 2238–2243.

Wolff, P., H. Jiang, G. Wang, J. Santos-Gonzalez and C. Kohler, 2015 Paternally expressed imprinted genes establish postzygotic hybridization barriers in Arabidopsis thaliana. eLife 4: 10.7554/eLife.10074.

Wu, C.-I., N. A. Johnson and M. F. Palopoli, 1996 Haldane’s rule and its legacy: why are there so many sterile males? Trends Ecol. Evol. 11: 281–284.

Zakaria, M., J. Ferent, I. Hristovska, Y. Laouarem, A. Zahaf et al., 2019 The Shh receptor Boc is important for myelin formation and repair. Development 146: dev172502.

Zechner, U., M. Reule, A. Orth, F. Bonhomme, B. Strack et al., 1996 An X-chromosome linked locus contributes to abnormal placental development in mouse interspecific hybrids. Nat. Genet. 12: 398–403.

Zechner, U., W. Shi, M. Hemberger, H. Himmelbauer, S. Otto et al., 2004 Divergent genetic and epigenetic post-zygotic isolation mechanisms in Mus and Peromyscus. J. Evol. Biol. 17: 453–460.

Zhang, B., and S. Horvath, 2005 A general framework for weighted gene co-expression network analysis. Stat. Appl. Genet. Mol. 4: 17.

Zou, H., D. Yu, X. Du, J. Wang, L. Chen et al., 2019 No imprinted XIST expression in pigs: biallelic XIST expression in early embryos and random X inactivation in placentas. Cell. Mol. Life Sci. 76: 4525–4538.

